# Error quantification in multi-parameter mapping facilitates robust estimation and enhanced group level sensitivity

**DOI:** 10.1101/2022.01.11.475846

**Authors:** Siawoosh Mohammadi, Tobias Streubel, Leonie Klock, Antoine Lutti, Kerrin Pine, Sandra Weber, Luke Edwards, Patrick Scheibe, Gabriel Ziegler, Jürgen Gallinat, Simone Kühn, Martina F. Callaghan, Nikolaus Weiskopf, Karsten Tabelow

## Abstract

Multi-Parameter Mapping (MPM) is a comprehensive quantitative neuroimaging protocol that enables estimation of four physical parameters (longitudinal and effective transverse relaxation rates *R*_*1*_ and 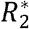, proton density *PD*, and magnetization transfer saturation *MT*_sat_) that are sensitive to microstructural tissue properties such as iron and myelin content. Their capability to reveal microstructural brain differences, however, is tightly bound to controlling random noise and artefacts (e.g. caused by head motion) in the signal. Here, we introduced a method to estimate the local error of *PD, R*_1_ and *MT*_sat_ maps that captures both noise and artefacts on a routine basis without requiring additional data. To investigate the method’s sensitivity to random noise, we calculated the model-based signal-to-noise ratio (mSNR) and showed in measurements and simulations that it correlated linearly with an experimental raw-image-based SNR map. We found that the mSNR varied with MPM protocols, magnetic field strength (3T vs. 7T) and MPM parameters: it halved from *PD* to *R*_1_ and decreased from *PD* to *MT*_sat_ by a factor of 3-4. Exploring the artefact-sensitivity of the error maps, we generated robust MPM parameters using two successive acquisitions of each contrast and the acquisition-specific errors to down-weight erroneous regions. The resulting robust MPM parameters showed reduced variability at the group level as compared to their single-repeat or averaged counterparts. The error and mSNR maps may better inform power-calculations by accounting for local data quality variations across measurements. Code to compute the mSNR maps and robustly combined MPM maps is available in the open-source hMRI toolbox.

## Introduction

Quantitative magnetic resonance imaging (qMRI) is more reproducible than conventional MRI (e.g. T1-weighted MRI typically used for morphometric analysis (Paus et al., 1999)) (Weiskopf et al., 2013; Cercignani and Bouyagoub, 2018). Quantification is typically achieved by acquiring multiple MRI contrasts to disentangle the mixture of different physical MR parameters present in conventional MRI, and correcting for instrumental variation through the acquisition of additional calibration measurements (Weiskopf et al., 2021). Multi-parameter mapping (MPM) provides a comprehensive approach to quantify multiple markers (such as longitudinal relaxation rate *R*_1_, proton density *PD*, effective transverse relaxation rate 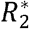, and magnetization transfer saturation *MT*_sat_) in a time-efficient MRI protocol composed of three multi-echo SPoiled Gradient Recalled echoes (SPGR) with PD-, T1-, and MT-weighting and additional calibration measurements (Helms et al., 2008a; Weiskopf et al., 2013) (Fig. 1). MPM parameters are sensitive to key biological microstructure features, e.g., myelin density and iron content (Kirilina et al., 2020), as well as volumetric changes. For example, the MPM parameters *R*_1_, *PD, MT*_sat_ sat have demonstrated utility in revealing aging processes (Callaghan et al., 2014), assessing clinical pathology (Freund et al., 2013; David et al., 2019) and illuminating behaviourally-relevant brain microstructure (Whitaker et al., 2016; Ziegler et al., 2019), and are known to be sensitive to macromolecular content and thus correlate with myelin density (West et al., 2018; Mohammadi and Callaghan, 2021). This sensitivity to microstructural tissue properties can be expected to vary between quantitative metrics depending on the underlying MR contrast mechanisms (Edwards et al., 2018; Weiskopf et al., 2021).

**Figure 1:**
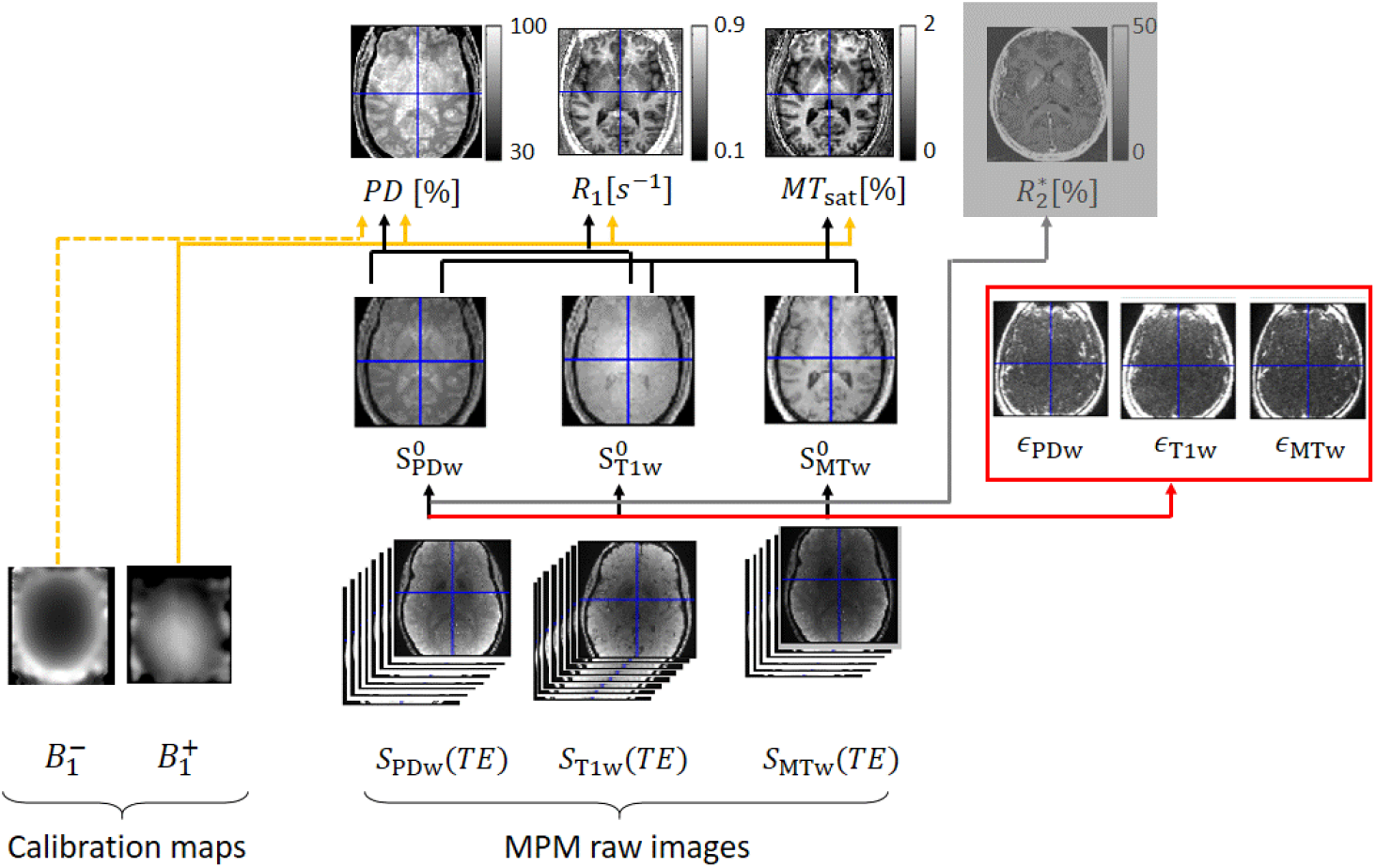
From raw data to parameter maps. Bottom row: the multi-parameter mapping (MPM) raw data as well as the receive 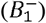 and transmit 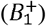 field maps acquired via dedicated calibration measurements. Middle row: the spoiled gradient-recall echo (SPGR) images with different contrasts at echo time (TE) zero (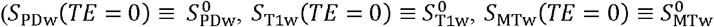, middle row) as well as the three contrast-specific uncertainties (*∈*_PDw_, *∈*_Tlw_, *∈*_MTw_, red box, bottom right), each of which summarizes the root mean-square difference between modelled and measured signal per contrast (*Background*, section 2). Top row: proton density (*PD*), longitudinal relaxation rate (*R*_1_), magnetization transfer saturation (*MT*_sat_), and mean-square difference between modelled and measured signal per contrast (*Background*, section 2). Top apparent transverse relaxation rate 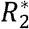, which is not considered here and thus greyed out. In the MPM framework (Tabelow et al., 2019), the three quantitative parameters *PD, R*_1_, and *MT*_sat_ are calculated from 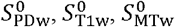 after correction for 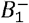 and 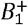 fields.

The comprehensive acquisition providing the four aforementioned quantitative parameters makes the MPM protocol particularly attractive for large-scale neuroimaging studies (Whitaker et al., 2016; Taubert et al., 2020; Clark et al., 2021) and clinical trials (Leutritz et al., 2020). When planning these kind of studies, a key question is whether we can objectively determine which metric will have the greater statistical sensitivity to the effect of interest under the influence of noise and artefacts (e.g. due to subject movement). In particular, clinical studies or those recruiting special populations depend on reliable power estimates to assess feasibility and efficiency. Power estimates and heuristics from traditional structural MRI techniques such as voxel-based morphometry (VBM) or other anatomical shape analyses cannot be translated to the analysis of quantitative MPMs, since they target largely different mechanisms (cluster of neighbouring voxels via the Jacobi-determinant modulation in VBM vs. single-voxel quantification in MPMs) and result in different characteristics for metrics such as scan-rescan reproducibility (Schnack et al., 2010). The sensitivity of the MPM parameters under the influence of random noise is often determined by the signal-to-noise ratio (SNR) of the underlying weighted volumes. Since we combine multiple weighted volumes with different SNRs to compute MPM parameters, the noise propagation into the MPM parameter estimates plays an important role for power estimation in MPM-based studies.

While sensitivity to microstructural brain differences promises the detection of more subtle anatomical effects than accessible by VBM, quantitative MRI maps are expected to be more susceptible to image artefacts than the constituent images. Moderate artefacts in the constituent images of quantitative MRI parameters can be amplified after nonlinear combination, e.g. when calculating the quantitative *PD, R*_1_ and *MT*_sat_ maps from the MPM raw data (Fig. 1). As a consequence, the variability of quantitative MRI parameters across a cohort is usually a composition of the true anatomical variability and variability due to biases caused by instrumental, physiological, and movement related outliers and noise (Weiskopf et al., 2014; Castella et al., 2018; Lutti et al., 2021). The outliers can significantly reduce the effective SNR of the data at the group level and thereby the sensitivity to microstructural changes. Thus, the MPM approach would greatly benefit from the quantification of parameter-specific errors that can routinely identify outliers on a voxel-wise basis.

In this study, we introduce a new method to estimate error maps for each of the three quantitative MPM parameters *R*_1_, *PD*, and *M*T_sat_ on a routine basis without the acquisition of any additional data (the error in R2* has been investigated elsewhere (Weiskopf et al., 2014)). The error maps are sensitive to two different types of variation: random noise, on the one side, and artefactual variation (e.g. due to imaging or subject motion), on the other. As a measure of the influence of random noise, we introduce the so-called model-based signal-to-noise ratio (mSNR) that is defined in analogy to the standard SNR, i.e. as the ratio between the MPM parameter and its error. First, we illustrate how the sensitivity to the two types of variation (random noise and artefacts) manifest themselves in the error and the mSNR maps. In two follow-up analyses, we investigate each of the two types of variation in more detail. The random-noise sensitivity is evaluated by quantifying the relation between the mSNR and the standard raw-image based SNR using both simulations and measurements. The artefact sensitivity is used to robustly combine MPM estimates from two successively acquired sets of MPM raw data (i.e. multi-echo SPGR images with PD-, T1-, and MT-weighting), where the acquisition-specific error maps are used to down-weight erroneous MPM parameters on a voxel-wise basis. We test the hypothesis that the robustly combined MPM estimates have lower image-artefact-related variability at the group level than the MPM estimates obtained from the arithmetic mean across acquisitions, or those obtained from a single MPM acquisition.

## Background

This section contains terminology and background information about the measurement of the signal-to-noise ratio (SNR) from raw data, the MPM estimation framework, and how the error (and by extension the model-based SNR) can be estimated within this framework. The raw-data SNR (Eq. (1)) is used as a reference to validate the utility of the mSNR and its capacity to estimate the per-acquisition SNR in the MPM parameters.

### Conventional SNR estimates of raw images

As a reference method to estimate SNR, we use the difference-image based SNR estimated from the raw multi-echo SPGR images with PD-weighting of a two-repeat measurement (Price et al., 1990; Reeder et al., 2005) or in short raw-image based SNR. For a twice repeated MPM measurement, the raw-image based SNR is estimated for the SPGR image with PD-weighting 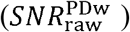 at the shortest echo time. The estimate of the mean signal is obtained from a small ROI by taking the mean of the signals from repeat 1 and 2, i.e. 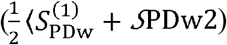, and this value is divided by the standard deviation of the difference of the signals from repeat 1 and 2 across the ROI, i.e. 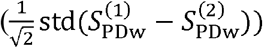, giving:

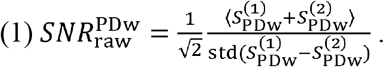

This method has been demonstrated to be a robust estimate of the SNR across different acceleration factors (Reeder et al., 2005; Dietrich et al., 2007). Note that an equivalent metric to that in Eq. (1) can also be calculated for the other two contrasts (MTw and T1w) and other echo times. Without loss of generality we focused on 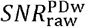 in this study.

### Error in MPM parameters

The noise level in the multi-echo SPGR images with PD-, T1-, and MT-weighting is estimated from the contrast-specific uncertainties (*ϵ*_PDw_, *ϵ*_T1w_, *ϵ*_MTw_ in Fig. 1) derived from the root-mean-square (rms) difference of the predicted and measured signal decay across echo times. The errors in the MPM parameters differ from the associated contrast-specific uncertainties of the PD-, T1-, and MT-weighted SPGR signal as illustrated in Fig. 2. This section summarizes how these metrics are related.

**Figure 2:**
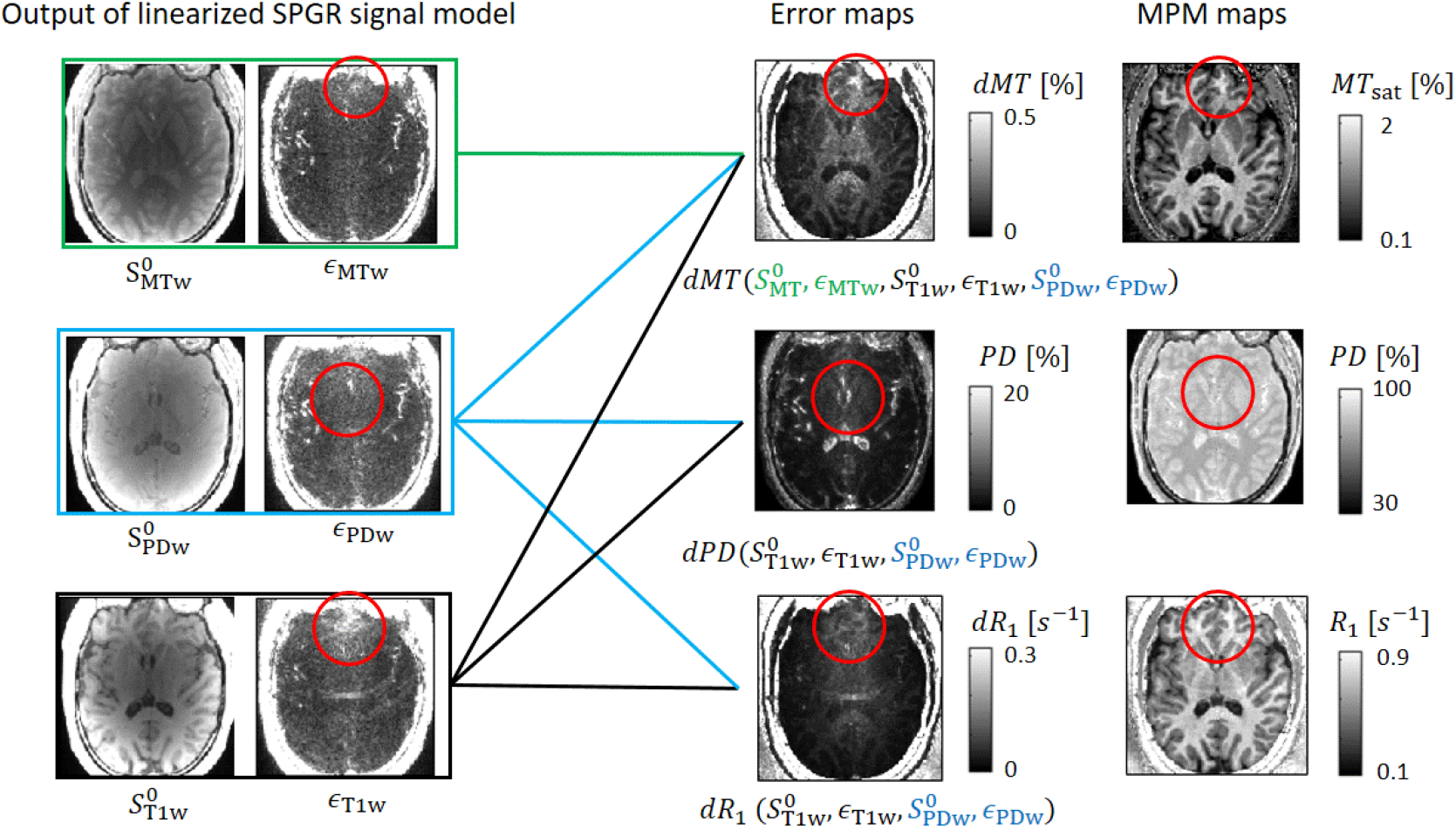
Error maps, their dependencies on MPM data and sensitivity to artefacts. Left column: the offset (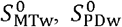, and 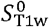) and contrast-specific uncertainties (*ϵ*_MTw_, *ϵ*_PDw_, and *ϵ*_Tlw_) of the SPGR signal as described in Figure 1. Middle column: the error maps and their dependencies (coloured lines) on offset and contrast-specific uncertainties. Right column: the MPM maps. The *MT* -error map (top-middle) depends on all offsets and contrast-specific uncertainties. In contrast, the *R*_1_-(bottom row) and *PT* -(middle row) error maps (d*R*_1_ and d*PD*), depend only on 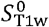 and 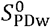 as well as on the corresponding contrast-specific uncertainties *ϵ*_Tlw_ and *ϵ*_PDw_. The red circles highlight regions with higher error or higher contrast-specific uncertainties and the corresponding region in the MPM parameters. Higher values in contrast-specific uncertainties are often smeared out (e.g. in *ϵ*_Tlw_) and localized increases are not necessarily accompanied by a biased MPM parameter (e.g. red circle in bottom row), whereas higher values in the error maps circle top row are co-localized with biased MPM parameters (e.g. circle in top row).

#### Error propagation

To estimate the error of each quantitative map, we calculated the first order propagation of error in *R*_1_, *PD*, and *MT*_sat_ under the assumption of uncorrelated errors between the SPGR and calibration measurements. For example, the error of *R*_1_ was derived to be:

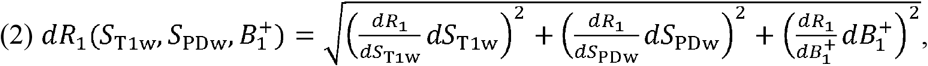

with, 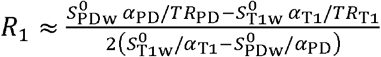. Local variations in the transmit field, 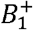 are incorporated into the flip angles via 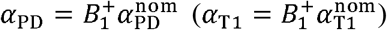 with 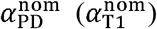 being the nominal flip angles (Helms et al., 2008a; Lee et al., 2017; Tabelow et al., 2019), 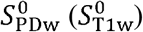 being the signal approximated at zero echo time *S*_PDw_ (*TE* = 0) *S*_T1w_ (*TE* = 0)) using the linearized SPGR signal fit (see next paragraph), and d*S*_R1_, and d*S*_PD_, being the contrast-specific uncertainties.

#### Linearized SPGR signal

To estimate the signal variation for each contrast, we first solved the joint model of the SPGR signals with PD-, T1-, and MT-weighting using the linearized exponential signal decay as introduced in the ESTATICS model (Weiskopf et al., 2014):

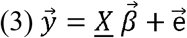

with 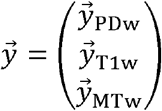 and 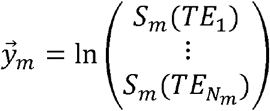, *<u>X</u>* being an N × 4 design matrix with rows *X*_*k,i*_ = [δ_PDw,*i*_, δ_T1w,*i*_, δ_MTw,*i*_, − *TE*_*k*_], *N*_*m*_ is the number of echoes for each contrast 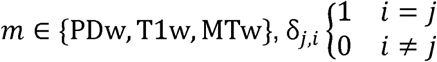 is the Kronecker delta, 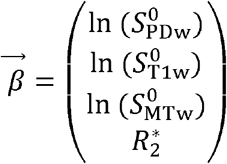 are regression coefficients, and the model-fit residual vector 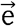 is composed of elements e_*k*_ that are equal to the difference between predicted and measured logarithmic signal for the k-th MPM volume *k* ∈ {1, … *N*_PDw_ + *N*_T1w_ + *N*_MTw_ }. While it has been previously shown that the elements of 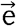 can be used to down-weight outliers for robust estimation of 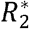 (Weiskopf et al., 2014), here, we use the rms difference of the predicted and measured signal as an estimate for the variation in the signal per contrast:

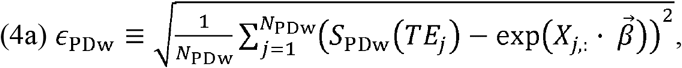

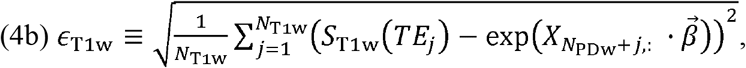

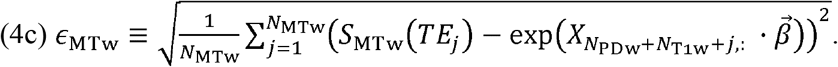

In the following, the measures in Eq. (4) are denoted as contrast-specific uncertainties (Fig. 1).

#### Error maps

Using the contrast-specific uncertainties (Eq. (4)) to characterise the noise within a given MPM acquisition, the error maps can now be estimated using the concept of error propagation. For the example of the error of R_1_ (Eq. (2)), the following is obtained:

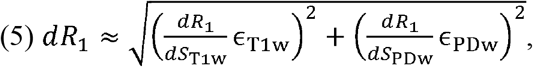

where the error in the transmit field is assumed to be negligible 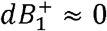. The same approach can be used to generate error maps for the *PD* and *MT*_sat_ estimates. Details of the derivation and the formulae of the errors for each parameter (*dR*_1_, *dPD, dMT*) can be found in Supplementary Materials 1-3 and their implementation in the hMRI toolbox can be found here: https://github.com/siawoosh/hMRI-toolbox.

#### Sensitivity of error maps to imaging artefacts

The relation between the error maps, the intercept of the linearized SPGR-signals which enters the error maps via the derivative of the MPM contrasts (e.g. for *dR*_1_ see the formulae for 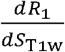 and 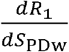 in the Supplementary Material, Eqs. [S3] and [S4]) and the contrast-specific uncertainties is illustrated in Fig. 2. We hypothesize that the error maps are sensitive to the factors that contribute to image degradation captured by the contrast-specific uncertainties (Eq. (4)), such as head motion during the acquisition of each image volume (Weiskopf et al., 2014; Castella et al., 2018), physiological artefacts, or sequence-specific artefacts (e.g. noise-enhancement due to parallel-imaging). Our hypothesis is motivated by the observation that artefacts in the contrast-specific uncertainty maps are often smeared out across the whole brain (e.g. in ϵ_T1w_, Fig. 2) and localized increases are not necessarily accompanied by a biased MPM parameter. High values in the error maps, however, are localized and coincide with noticeable variation in the associated MPM parameter maps (red circles in top row, Fig. 2).

### Model-based SNR (mSNR)

Following the definition of SNR, we introduce here the model-based SNR maps (in short mSNR maps) for each MPM parameter map. This is defined as the ratio between a quantitative MPM map and its corresponding error map. For example, the mSNR map for the *R*_1_ parameter was calculated as follows:

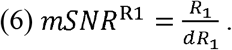

To avoid divergence for very small error values, a threshold was used to set the mSNR map to zero when dR1 < 10^−4^ (the same was done for *mSNR*^MT^ and *mSNR*^PD^, with dMT < 10^−4^ and dPD <^−2^ respectively). These thresholds were chosen heuristically. Note that the MPM-specific mSNRs are estimated per MPM acquisition.

### Robust combination of MPM parameters

When multiple MPM datasets are available, the resulting parameters can be combined by a simple arithmetic mean. Here we propose an alternative robust combination, of potentially erroneous MPM parameters, by exploiting their empirical error maps. The idea is formulated for the case of two distinct imaging repeats but can be generalized to multiple repeats.

To generate robust MPM parameters (denoted by superscript “RO”) from a two-repeat protocol a function of the error maps is used to weight each repeat according to their voxel-wise error. For the example of *R*_1_ the robust-combination was defined as follows on a voxel-by-voxel basis:

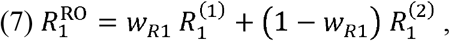

where. 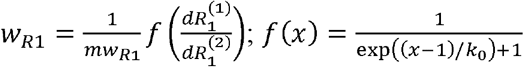 is the Fermi function; 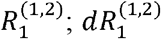 are the longitudinal relaxation rates and their respective errors from repeats (1) and (2), all defined on a per voxel basis; and. 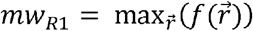 is defined as the maximum weight across voxels 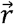. The parameter *k*_0_ tunes the sensitivity of the weights with respect to the ratio of errors: small *k*_0_ leads to high sensitivity. The parameter (*k*_0_ = 0.1) was heuristically optimized for one subject that showed strong motion artefacts in the scan-rescan measurements (see Figs. 5-7) and applied to the rest of the subjects. This parameter can be adjusted for different protocols via the local default file of the hMRI toolbox.

## Materials and Methods

### Subjects and MRI

#### Subjects

20 healthy volunteers participated in this study. 18 were measured with protocol 1, one with protocol 2 at 3T, and one with protocol 3 at 7T. We excluded two participants measured with protocol 1 from the analysis: one due to excessive, unsalvageable levels of movement and one due to image reconstruction problems. We include”d th e remaining 16 participants in the reported group analysis (age: 20-54 years; *mean*(*SD*) = 32.63(8.55) years; 7 female, 9 male). Exclusion criteria were any psychiatric disorders, assessed via the Mini-International Neuropsychiatric Interview (Ackenheil et al., 1999), neurological diseases, head trauma or metallic implants. Participants provided written informed consent and were compensated for their participation. The local ethics committees at University Medical Center Hamburg-Eppendorf and Medical Faculty of the University of Leipzig approved the study (PV5141; LPEK_006_Kühn; Reg.-No. 273-14-25082014).

#### MRI protocol

Scans were performed on three MRI systems (Siemens Healthineers, Erlangen, Germany): 3T PRISMA (protocol 1), 3T PRISMA-fit (protocol 2), and Magnetom 7T (protocol 3). For protocols 1 and 2, the body coil was used for transmission (Tx) and the 64-channel receiver head-coil for reception (Rx). For protocol 3, an integrated 1-channel Tx/32-channel Rx head coil (Nova Medical, Wilmington, MA, USA) was used. Whole brain MR images were acquired using the MPM (Weiskopf et al., 2013) protocol, including three differently weighted (MT-, PD- and, *T*_1_-weightings) multi-echo SPGR contrasts (protocol 3 only acquired two weightings, PD- and, *T*_1_-weighting). Rapid calibration data were acquired at the outset of each repeat to correct for inhomogeneities in the RF transmit field (Lutti et al., 2010, 2012) including a B0 field map for correction of EPI distortions in the transmit field map. For protocols 1 and 2, the automatic transmit adjust procedure was used, whereas for protocol 3, the transmit voltage was calibrated using an initial low-resolution transmit field map to be optimal over the occipital lobe.

The sequence parameters for protocols 1-3 are summarized in Table 1.

**Table 1:**
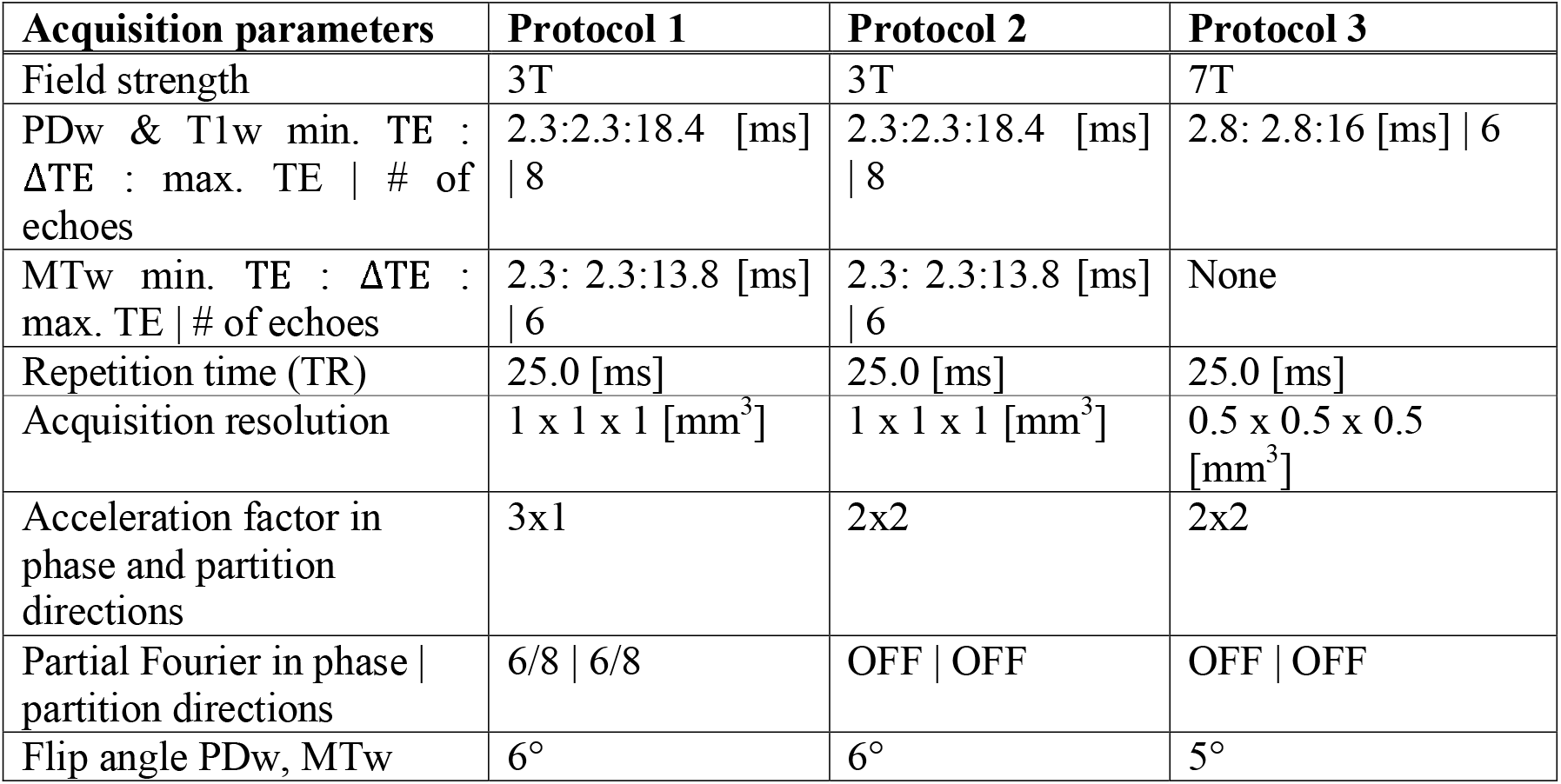
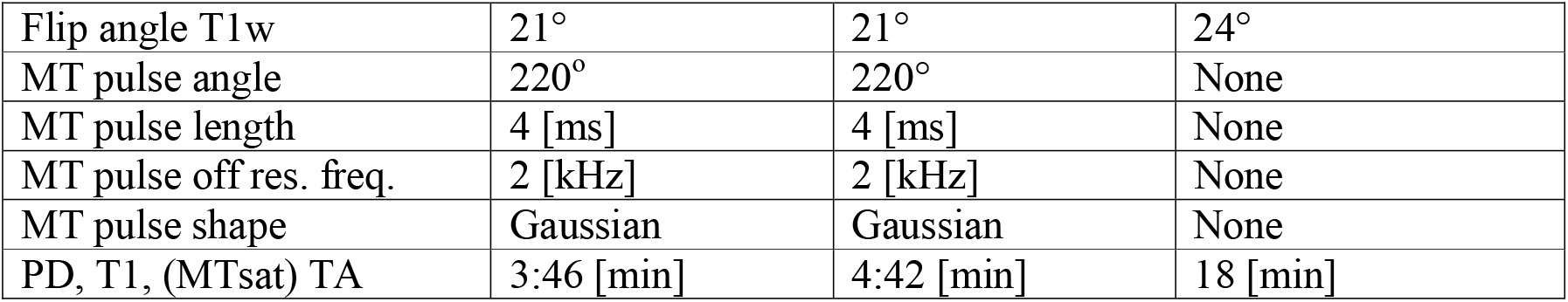
MRI parameters. Different sequence parameters for protocols 1-3. Note that protocol 3 does not include an MTsat map. Abbreviations: FoV, field of view, MT(w), magnetization transfer (weighted), PD(w), proton density (weighted), T1w, T1 weighted, T1, longitudinal relaxation time, TE, echo time, TA, acquisition time per set of multi-echo SPGR images, SPGR, spoiled gradient-recall echo.

| Acquisition parameters | Protocol 1 | Protocol 2 | Protocol 3 |
| --- | --- | --- | --- |
| Field strength | 3T | 3T | 7T |
| PDw & T1w min. TE : ΔTE : max. TE \| # of echoes | 2.3:2.3:18.4 [ms] \| 8 | 2.3:2.3:18.4 [ms] \| 8 | 2.8: 2.8:16 [ms] \| 6 |
| MTw min. TE : ΔTE : max. TE \| # of echoes | 2.3: 2.3:13.8 [ms] \| 6 | 2.3: 2.3:13.8 [ms] \| 6 | None |
| Repetition time (TR) | 25.0 [ms] | 25.0 [ms] | 25.0 [ms] |
| Acquisition resolution | 1 x 1 x 1 [mm<sup>3</sup>] | 1 x 1 x 1 [mm<sup>3</sup>] | 0.5 x 0.5 x 0.5 [mm<sup>3</sup>] |
| Acceleration factor in phase and partition directions | 3x1 | 2x2 | 2x2 |
| Partial Fourier in phase \| partition directions | 6/8 \| 6/8 | OFF \| OFF | OFF \| OFF |
| Flip angle PDw, MTw | 6° | 6° | 5° |
| Flip angle T1w | 21° | 21° | 24° |
| MT pulse angle | 220° | 220° | None |
| MT pulse length | 4 [ms] | 4 [ms] | None |
| MT pulse off res. freq. | 2 [kHz] | 2 [kHz] | None |
| MT pulse shape | Gaussian | Gaussian | None |
| PD, T1, (MTsat) TA | 3:46 [min] | 4:42 [min] | 18 [min] |

The acquisition of all multi-echo SPGR contrasts was repeated for each individual. This was done within a single imaging session at 3T (i.e. protocols 1 and 2 contained two “runs” of each contrast) and in two separate imaging sessions at 7T (i.e. protocol 3 contained only one run).

The total scan time of both runs was about 28 min (=2×11+6) for protocol 1, about 33 min (=2×13.5+6) for protocol 2, and about 84min (=2x(36+6)) min for protocol 3.

For protocol 3, participant motion was monitored and corrected prospectively by an optical tracking system (Kineticor, Honolulu, HI, USA) (Callaghan et al., 2015). Each volunteer was scanned while wearing a mouth guard with a passive Moiré pattern marker used for tracking (manufactured by the Department of Cardiology, Endodontology and Periodontology, University Medical Center Leipzig; comparable to (Papoutsi et al., 2020)). The dataset acquired with protocol 3 was taken from the study by (McColgan et al., 2021).

In all protocols, parallel imaging was performed using generalised autocalibrating partial parallel acquisition (GRAPPA) (Griswold et al., 2002) with acceleration factors of 3 or 4.

### Map creation and spatial processing

Map creation and spatial processing was performed using modules in SPM12 version v7771 (Friston et al., 2006) and a branch of the hMRI toolbox (Tabelow et al., 2019) available here: https://github.com/siawoosh/hMRI-toolbox.

#### Rigid-body registration

To ensure that the data from both repeats were in the same space, the second dataset was registered to the first using a rigid-body transformation (spm_coreg). To this end, first the MPM maps were aligned to the MNI-space template (“avg152T1”) with the auto-align module in the hMRI toolbox (Tabelow et al., 2019) using the *MT*_sat_ map as “source image” (i.e. the image that was used for estimating the transformation parameters). Then, the *MT*_sat_ maps were thresholded (*MT*_sat_>0 and *MT*_sat_<5 p.u.) to remove unreasonable or extreme values and segmented (spm_segment) into grey and white matter tissue probability maps (TPMs). These grey and white matter TPMs were combined and used as source and target images for inter-repeat registration, with the TPMs from the second repeat being the source and the TPMs of the first repeat being the target image. The estimated transformation parameters were applied to all maps from the second repeat. The reason for using the grey and white matter TPMs for registration instead of the original maps was to reduce potential confounding effects of motion artefacts in the individual MPM images on the accuracy of the registration.

#### Non-linear spatial registration and spatial smoothing in common group space

First the MPM maps of each subject were transformed into MNI space using the geodesic shooting nonlinear registration tools (spm_shoot, (Ashburner and Friston, 2011)). To estimate the nonlinear transformation that maps each individual brain into common space, high-quality grey and white matter tissue probability maps were generated per subject by segmenting the arithmetic mean of the *MT*_sat_ maps across both repeats. Then, the hMRI toolbox was used for tissue-specific smoothing (Tabelow et al., 2019).

### Analysis

Three analyses were performed, which we describe in detail below. In the first analysis, we assessed two different features of the error and mSNR maps: their sensitivity to the image SNR and to image artefacts. Second, the relation between mSNR and the image noise of SPGR raw images was quantitatively investigated. Third, the artefact-associated variability of MPM parameters at the group level was investigated for different combinations of a two repeat MPM protocol.

#### Analysis I: Illustration of error and mSNR maps

On the single-subject level, we investigated how the mSNR varied for different MPM protocols and how imaging artefacts in the MPM parameter maps appeared in the error and the mSNR maps. For the same subject, we investigated in two repeats with varying artefact levels the correspondence between error and respective MPM parameter maps in erroneous regions and how the MPM parameter values per repeat could be weighted towards the less erroneous repeat in the robust combination.

#### Analysis II: Quantifying relation between raw-image-based and model based SNR

The propagation of SNR into mSNR was characterized by simulations and in vivo measurements. Hereby, we used the raw-image-based SNR of the PD-weighted image (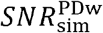, see Eq. (1)) as a proxy for the image SNR, both in the simulations and measurements. To assess the dependence between *SNR*^PDw^ and *mSNR*^*m*^ for varying SNRs, a linear model was fit to the data:

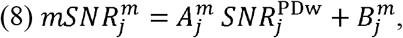

with 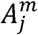 and 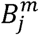 being the fitted parameters, *m* ∈ {PD, R1, MT} and *j* being the index that specifies whether simulated or measured data was used (*j* ∈ {sim,meas}).

#### Simulated 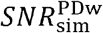 and 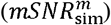

To simulate the noisy SPGR signal, we added complex-valued Gaussian noise to the rational approximation of the Ernst equation (Ernst and Anderson, 1966; Helms et al., 2008a) with heuristic approximation of the MT-pulse effect (Helms et al., 2008b). Then, the absolute value of the noisy signal was calculated as follows:

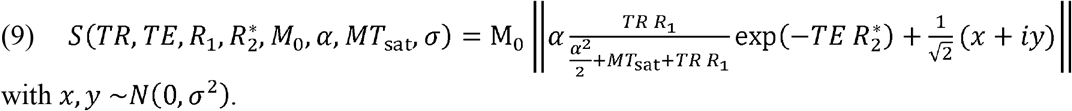

The multi-echo SPGR signals were simulated using Eq. (9) and the parameters in Table 2. To achieve PD- and T1-weighting using the respective flip angles for protocol 1 (see Table 1) and, *MT*sat = 0. The MT-weighted multi-echo SPGR signal was simulated using the same parameters as for the PD-weighted signal only with the difference that the *MT*_sat_ value in Table 2 was used. Then, *mSNR*^PD^, *mSNR*^R1^, and *mSNR*^MT^, were calculated from the simulated SPGR signals using the proposed approach (see Background and Supplementary Material 1-3). Additionally, 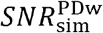 was calculated from the simulated signal using Eq. (1). Note that the simulation was performed separately for white matter and grey matter ground truth parameters (Table 2) to investigate the influence of tissue type on the relation between 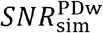 and *mSNR*^m^.

**Table 2:**
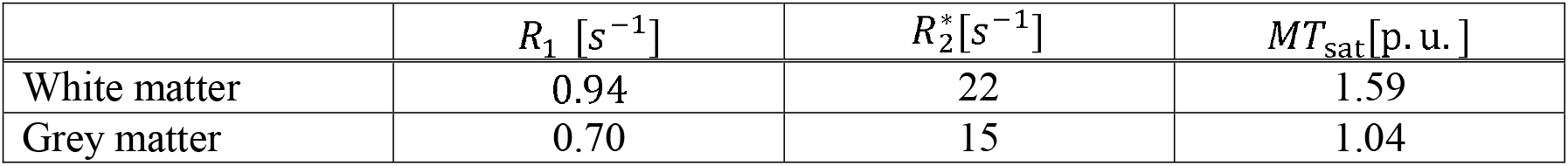
Simulation parameters. These parameters were used to simulate the signal in Eq. (9) for grey and white matter. Additionally, the net magnetization was set to 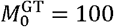 and other parameters (repetition time: *TR*, echo time: *TE*) were as in protocol 1. Finally, the zero-mean, additive Gaussian noise *N*(0,*σ*^2^) with varying standard deviation was added (*σ* ∈ {0.002,…,0.1}), each time with 5000 noise realisations. This resulted in 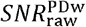 values between 62 and 2.

#### Measured 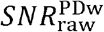 and mSNR^m^

To partition the MRI brain volumes into regions of interest (ROIs) with varying SNR, we used the Oxford-Harvard atlas for grey and white matter (Frazier et al., 2005; Desikan et al., 2006; Makris et al., 2006; Goldstein et al., 2007). Hereby, it was assumed that regions closer to the skull have higher SNR because they are closer to the head coil than regions within the centre of the brain. Note that this analysis assumes that the proposed SNR measures are independent of the tissue type (i.e. whether it is grey or white matter) but solely depend on their distance to the head coil.

The SNR measures were estimated for each subject in individual space, to prevent interpolation artefacts associated with the spatially non-linear registration into common space affecting the estimation of 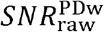. To this end, ROIs were projected into individual space using the inverse of the spatial transformations estimated in the “spatial processing section”. Then, the individual tissue probability maps for white and grey matter were multiplied with each individual ROI and thresholded at 90%. Only ROIs containing more than 100 voxels were used for the analysis. Then, 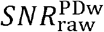 and mSNR^m^ were estimated within each ROI for each subject. This experiment was performed for the data acquired with protocol 1.

Additionally, the subject-averaged 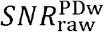 per ROI was projected into a template image in group space for visualization purposes only. The spatial dependence of mSNR maps was qualitatively compared for two different protocols at 3T (protocol 1 and 2) and one at 7T (protocol 3). Then, the average mSNR value across the brain was quantified per MPM parameter and protocol (averaged across subjects for protocol 1).

#### Analysis III: Artefactual variation at the group level

The variability of MPM parameters across a group of healthy subjects is expected to be a combination of the true anatomical variability in the cohort and artefactual variability caused by instrumental, physiological, and subject-movement related noise. While the latter can be reduced by averaging, the former cannot. To assess the variability across the group, we calculated the standard-error-of-the-mean (SEM) for four sets of MPM parameters in MNI space after tissue-specific hMRI smoothing (see section “spatial processing”). The four sets of MPM parameters were generated either from repeats (1) and (2) separately, or from the arithmetic mean (AM) across repeats or their robust combination (RO in *Eq*. (7)).

To assess the effect of instrumental, physiological, and subject-movement on the variability, the SEM of the arithmetic-mean combined MPM parameters 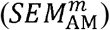 were used as reference (with *m* ∈ {MT, PD, *R*1}) and compared to the SEM of the other sets of MPM parameters via their relative difference: 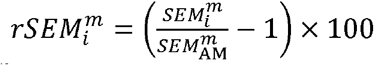 with *i*. ∈{1,2 RO}. Hereby, a negative (positive) 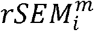 would indicate that the variability in the *I* dataset is smaller (larger) than in the reference AM dataset. Since it is expected that instrumental, physiological, and subject-movement artefacts increase the variability of the estimated parameters, negative 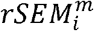 values in each dataset were interpreted as a reduction of artefactual variability and thus an increase of sensitivity towards group differences whereas positive 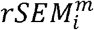 values were interpreted as a decline of sensitivity.

## Results

### Analysis I: Illustration of error and mSNR maps

This analysis illustrates how variations due to image SNR and artefacts manifest themselves in the mSNR and error maps for different MPM protocols and at different field strengths. The different protocols showed a decrease of mSNR towards the centre of the brain. Although protocols 1 and 2 were measured with similar parameters (e.g. spatial resolution, TE, TR) and instruments (e.g. at 3T and with a 64ch head-coil), the decrease towards the centre of the brain was differently shaped within the respective mSNR maps: the mSNR maps measured with protocol 1 had an elliptical pattern of lower mSNR centrally, the long axis of which ran anterior-posterior whereas the central region of lower mSNR was circular for protocol 2, but decreased less rapidly than protocol 1 (dashed lines in Fig. 3a). The decline of mSNR was accompanied by an increase in the average contrast-specific uncertainty for protocol 1 (1^st^ row, 4^th^ column, Fig. 3a), indicating that the noise or artefact-level is higher in the areas of decreased mSNR. Although *mSNR*^PD^, *mSNR*^R1^ maps measured with protocol 3 showed smaller gradients towards the centre of the brain as compared to their counterparts measured with protocols 1 and 2, they revealed the same trend (higher mSNR values towards the cortex and lower towards the centre of the brain). On average, across the brain, we found that the mSNR values decreased from protocol 1 to 3 (Fig. 3b): the mSNR of protocol 1 was about 1.2 times larger than the mSNR of protocol 2, whereas the ratio between the mSNR values of protocols 1 and 3 was about 1.6 for *PD* and *R*_1_. The ratio of *mSNR*^PD^ to *mSNR*^R1^ was protocols 1 and 3 was about 1.6 for *PD* and *R*_1_. The ratio of *mSNR*^PD^ to *mSNR*^R1^ was similar across protocols: 1.79 for protocol 1, 1.80 for protocol 2, and 1.82 for protocol 3.

**Figure 3:**
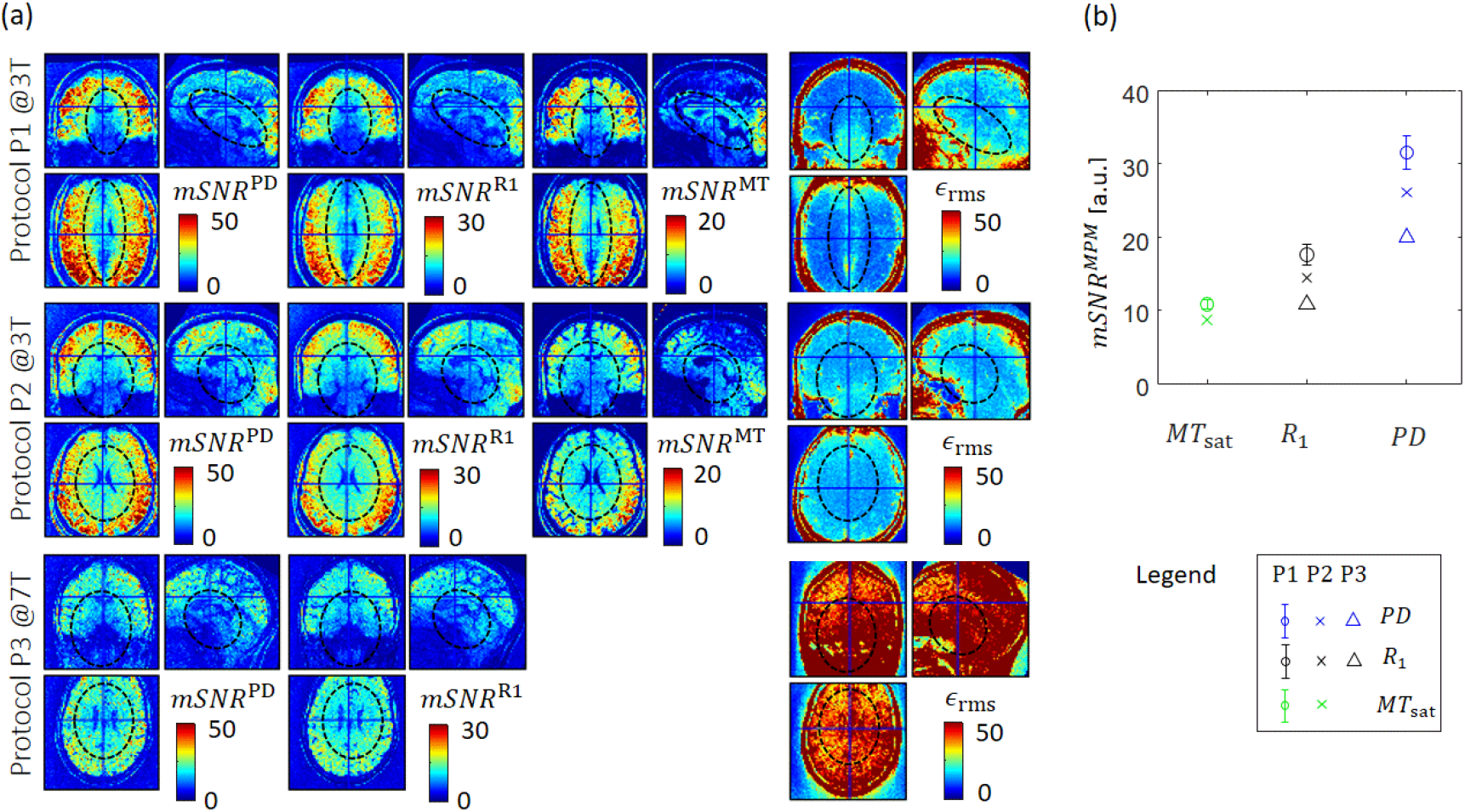
Variation of mSNR maps across the brain for different MPM protocols. (a): mSNR maps for three MPM protocols (P1-P3) are depicted (1^st^ row: P1, 2^nd^ row: P2, and 3^rd^ row: P3): *mSNR*^PD^ (1^st^ column), *mSNR*^R1^ (2^nd^ column), *mSNR*^MD^ (3^rd^ column). Fourth column: the averaged across contrast-specific uncertainties is depicted (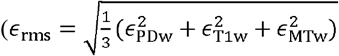, 4^th^ column). (b) The mSNR parameters averaged across the brain for protocols P1 (circle), P2 (cross), and P3 (triangle) for the three MPM parameters (*PD*: blue, *R*_l_: black, *MT*_sat_: green). Note that P1 was calculated across a group of healthy subjects (standard deviation across group in black) whereas P2 and P3 were calculated only for a single subject (no standard deviation). *mSNR*^m^ and *ϵ*_rms_ in arbitrary units. The values in the mSNR maps decrease towards the centre of the brain accompanied by an increase in the rms values. While the mSNR map measured with protocol 1 showed a strong left-right gradient (middle row, left: ellipse with large eccentricity), the mSNR maps measured with protocols 2 (and two out of three mSNR maps measured with protocol 3) showed a circular shaped area of decreased values (middle row: middle and right). Note that, since no MT measurement was available for protocol 3, *ϵ*_rms_ was the mean of the remaining two residuals.

Fig. 4 shows the relation between each MPM parameter, the associated error and mSNR maps. The mSNR maps had the same contrast for all MPM parameters, while the contrast in the error maps reflects the MPM parameter contrast (*PD*: high in grey matter and low in white matter, *MT*_sat_ and *R*_1_ : low in grey matter and high in white matter).

**Figure 4:**
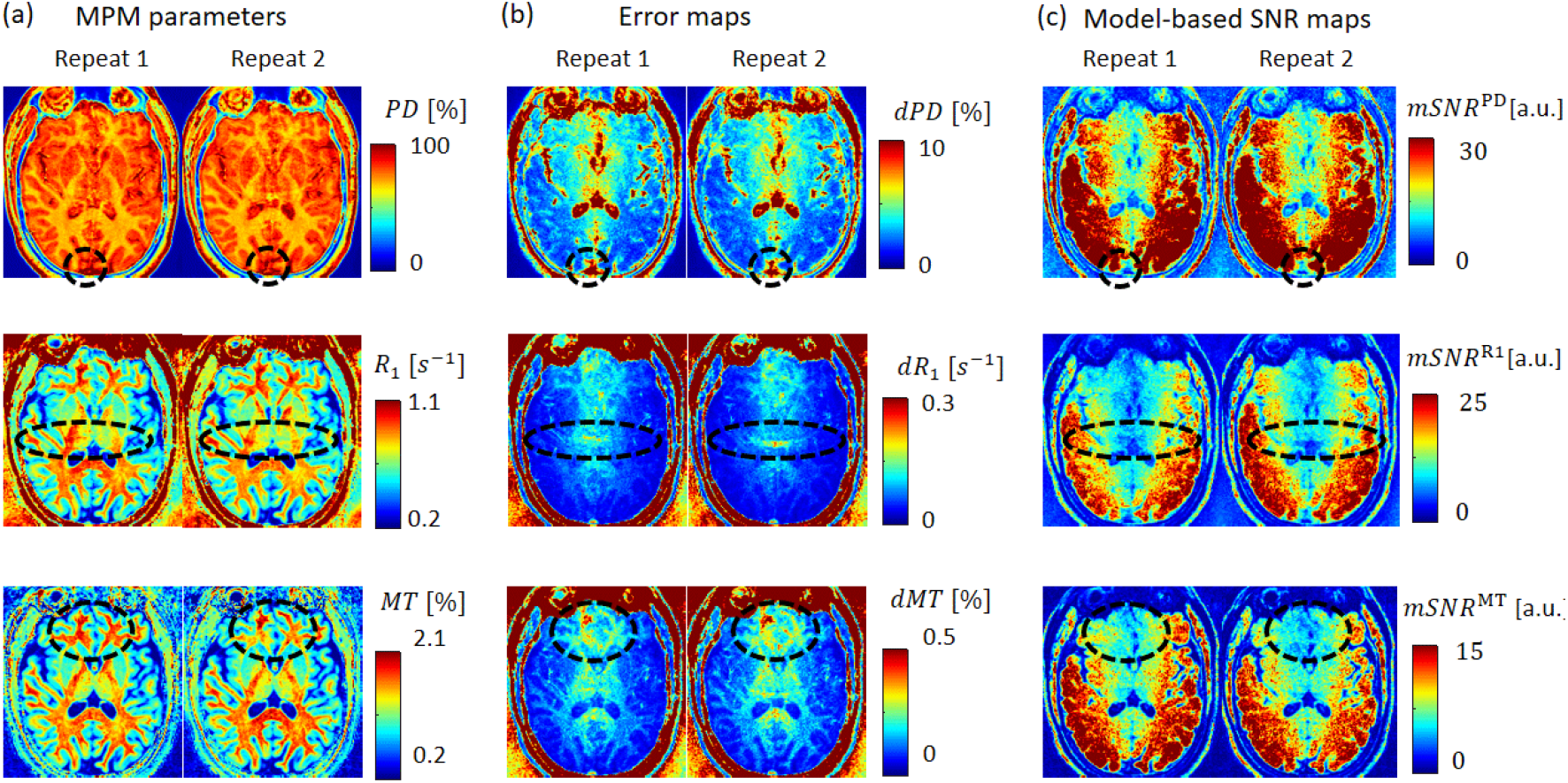
Artefacts in the MPM, error, and mSNR maps. Depicted are the quantitative MPM parameters (left column), the error maps (middle column), and corresponding mSNR maps (right column) for a subject, measured in two repeats with varying artefacts using protocol 1. The mSNR maps have the same contrast, even though the contrast of the associated MPM map varies (top: *PD*, top: *R*_1_, bottom: *MT*_sat_). The artefacts in the parameter maps (highlighted by dashed circles) manifest themselves as increased error-map values and reduced mSNR-map values. Three artefacts were identified with potentially different origin: physiological noise that remained almost the same between repeats at the superior sagittal sinus (top row), aliasing artefacts that varied between repeats (middle row), and voluntary subject motion artefacts that strongly varied between repeats (bottom row). Note that intensity ranges for the three mSNR maps differ.

As expected, artefacts in the MPM maps (highlighted by dashed circles) manifested differently in the error and mSNR maps: while the error in regions affected by artefacts was increased, the mSNR value was decreased because the latter depends reciprocally on the error. Three artefactual regions with potentially different artefact-causes were identified. The artefact in the top row may be caused by physiological noise due to flow artefacts because it was located in the superior sagittal sinus and did not vary between repeats. In the middle row, an aliasing artefact was identified which varied between repeats and thus might be enhanced by involuntary subject motion. The artefact highlighted in the bottom row was most likely a result of involuntary subject motion because it varied between the two repeats.

Figures 5-7 illustrate regionally localized artefacts in the MPM parameters that were captured by the error maps, became less pronounced in the arithmetic mean and could be partly removed in the robust combination (Fig. 5 for *PD*, Fig. 6 for *R*_1_, and Fig. 7 for *MT*_sat_). These artefacts were probably related to involuntary subject-motion because they varied between repeats.

**Figure 5:**
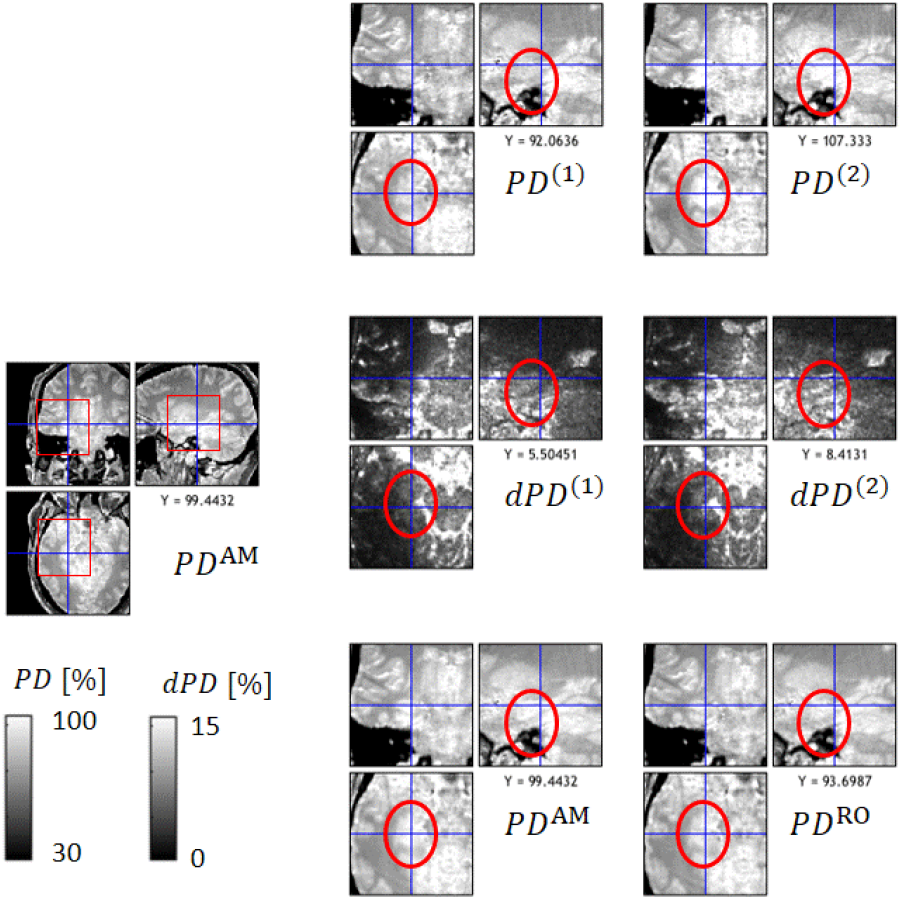
Reduced artefacts in robustly-combined proton density (*PD*) map. Depicted are: two successive repeats of the PD map using protocol 1 with superscript *(1)* and *(2)* (top row), the associated error maps for each repeat (middle row), and their arithmetic mean and robustly combined average with superscript *AM* and *RO* (bottom row). An area is magnified (red box, left column), where the error maps were sensitive to artefacts (hyper intensities) and the robust combined PD contained less artificially increased values than the arithmetic mean (circle) and single-repeat PD maps.

**Figure 6:**
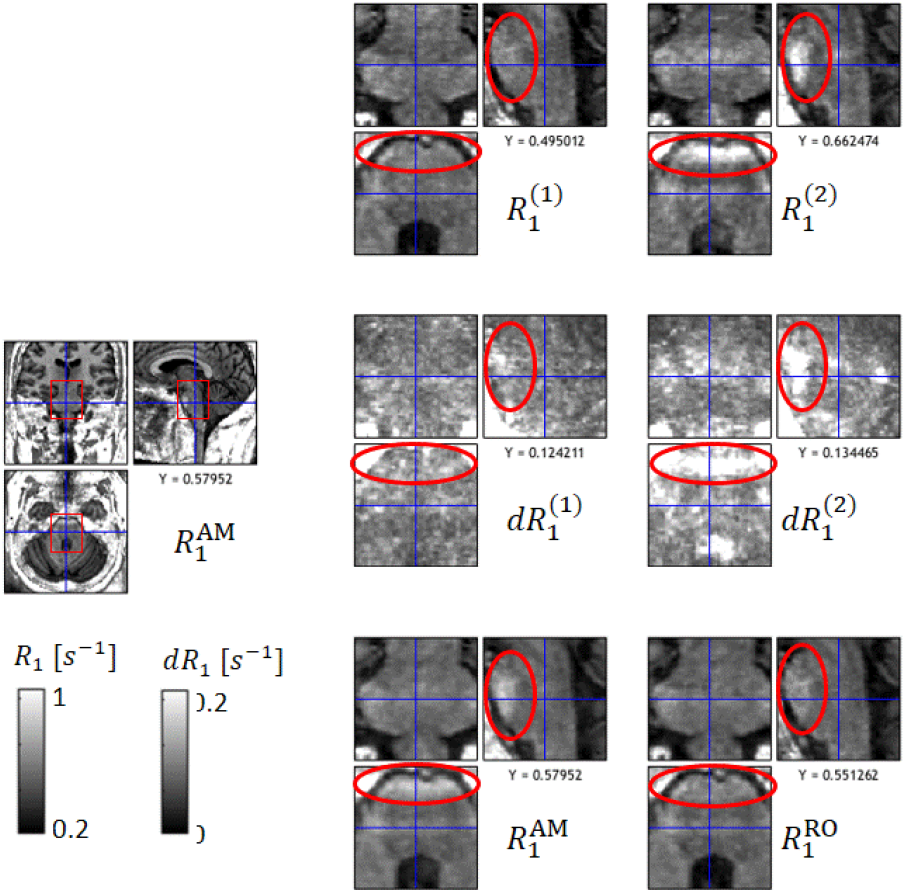
Reduced artefacts in robustly-combined longitudinal relaxation rate (*R*_1_) parameter. Depicted is the same information as in Figure 5 for the *R*_1_ parameter instead of the *PD* parameter.

**Figure 7:**
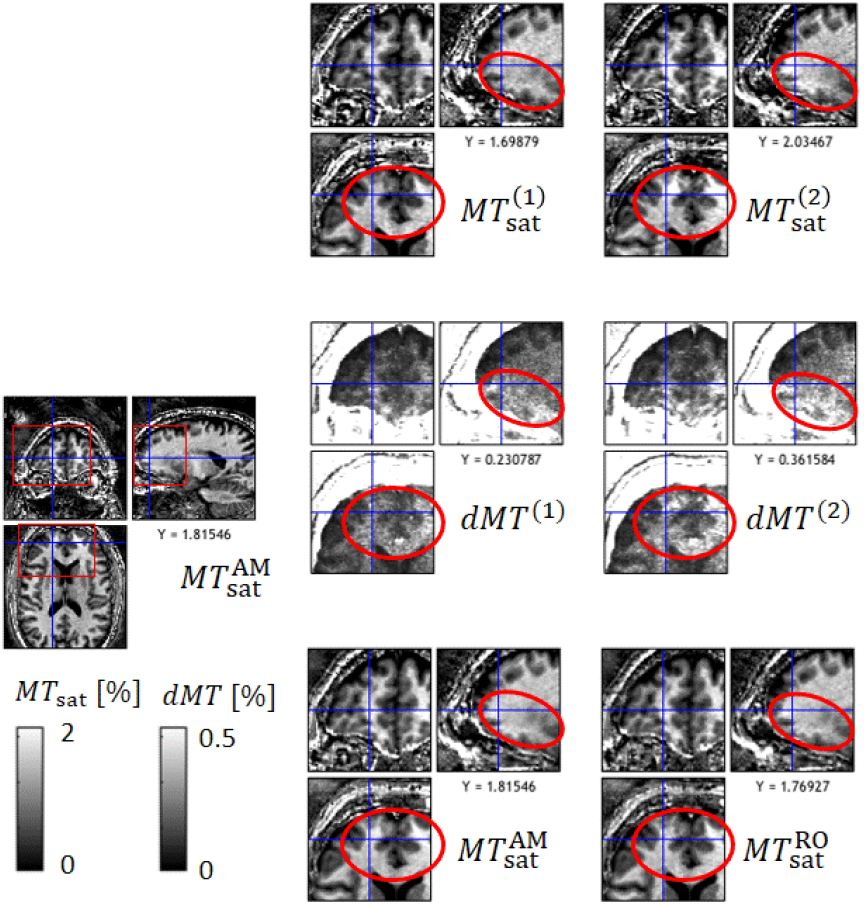
Reduced artefacts in robustly-combined magnetization transfer saturation rate (*MT*_sat_) parameter. Depicted is the same information as in Figure 5 for the *MT*_sat_ parameter instead of the *PD* parameter.

### Analysis II: Quantifying relation between raw-image and model based SNR

Here, we quantify a linear relation between image SNR and mSNR at the group level and in simulations. Fig. 8 depicts the linear relation between 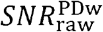 and *mSNR*^*m*^ of the three MPM parameters (*m* ∈ {PD,R1,MT}) in simulation (Fig. 8a-c) and in measurements (Fig. 8d-f) with slopes and intercepts reported in Table 3. For the measurement, variation in 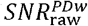 and *mSNR*^m^ was achieved by measuring the respective metrics within 111 regions of interest (ROIs) averaged across 16 healthy subjects using protocol 1 (Fig. 9).

**Table 3:**
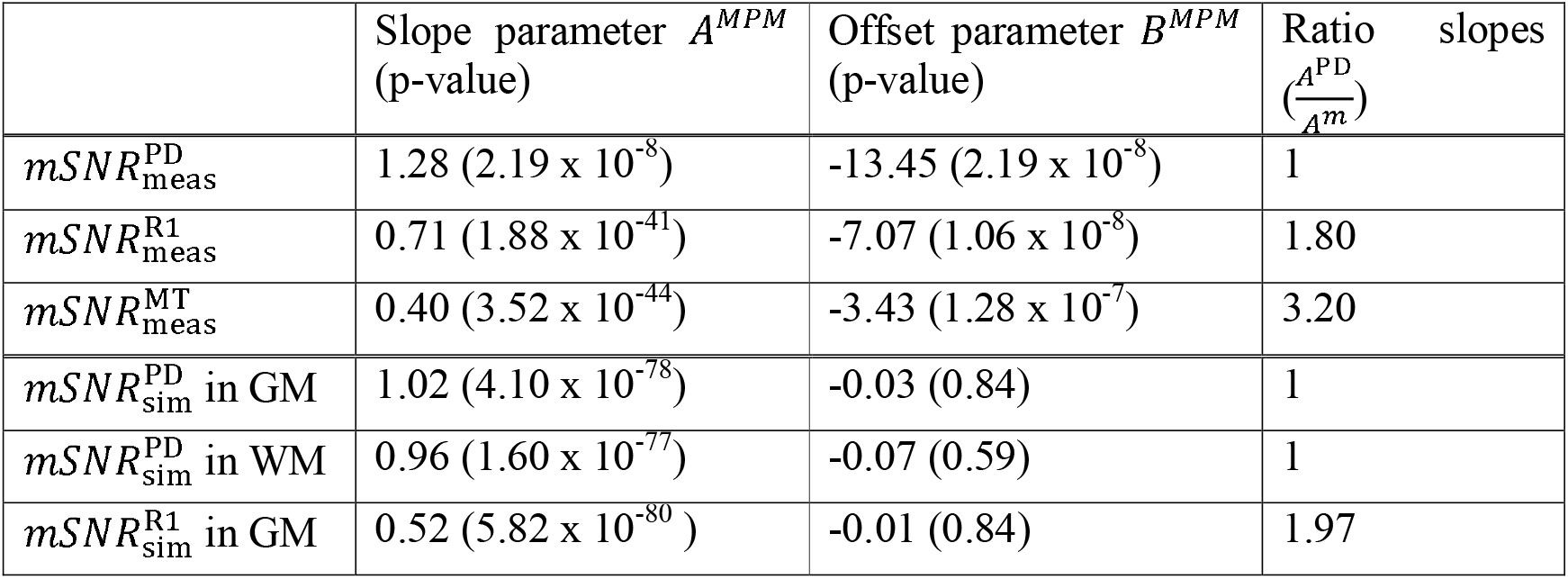
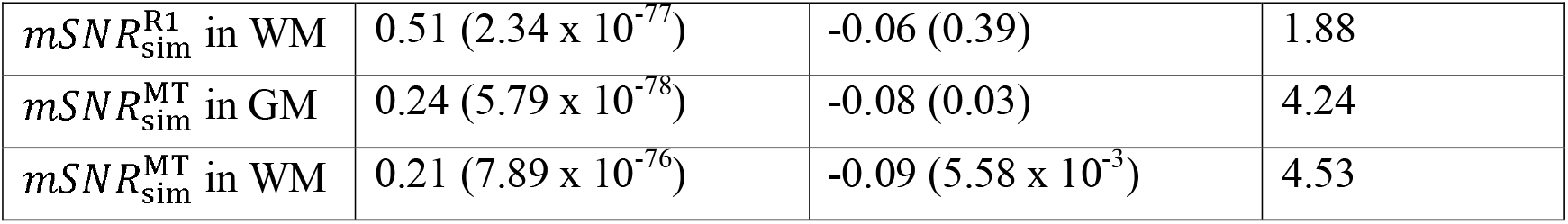
Simulated and measured relation between model-based and raw image-based SNR. The coefficients of the heuristic linear models (Eq. (8)) that relate the 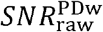 to the simulated 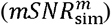 and measured 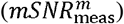 mSNR, summarizing the slopes and intercepts of the curves in Figure 8, with *A*^*m*^ being the slope and *B*^*m*^ being the intercept, and the ratio between *A*^PD^ and *A*^*m*^ (*m* ∈ {*PD*,R1,MT}). The subscript “sim” refers to the simulated data in Figure 8a-c and “meas” to the measured data in Figure 8e-f (for details see methods section “Analysis II”). Note that for the measured mSNR the average across repeats was used.

| | Slope parameter $A^{MPM}$<br>(p-value) | Offset parameter $B^{MPM}$<br>(p-value) | Ratio slopes<br>$(\frac{A^{PD}}{A^m})$ |
| --- | --- | --- | --- |
| $mSNR_{meas}^{PD}$ | 1.28 ($2.19 \times 10^{-8}$) | -13.45 ($2.19 \times 10^{-8}$) | 1 |
| $mSNR_{meas}^{R1}$ | 0.71 ($1.88 \times 10^{-41}$) | -7.07 ($1.06 \times 10^{-8}$) | 1.80 |
| $mSNR_{meas}^{MT}$ | 0.40 ($3.52 \times 10^{-44}$) | -3.43 ($1.28 \times 10^{-7}$) | 3.20 |
| $mSNR_{sim}^{PD}$ in GM | 1.02 ($4.10 \times 10^{-78}$) | -0.03 (0.84) | 1 |
| $mSNR_{sim}^{PD}$ in WM | 0.96 ($1.60 \times 10^{-77}$) | -0.07 (0.59) | 1 |
| $mSNR_{sim}^{R1}$ in GM | 0.52 ($5.82 \times 10^{-80}$) | -0.01 (0.84) | 1.97 |
| $mSNR_{sim}^{R1}$ in WM | 0.51 ($2.34 \times 10^{-77}$) | -0.06 (0.39) | 1.88 |
| $mSNR_{sim}^{MT}$ in GM | 0.24 ($5.79 \times 10^{-78}$) | -0.08 (0.03) | 4.24 |
| $mSNR_{sim}^{MT}$ in WM | 0.21 ($7.89 \times 10^{-76}$) | -0.09 ($5.58 \times 10^{-3}$) | 4.53 |

**Figure 8:**
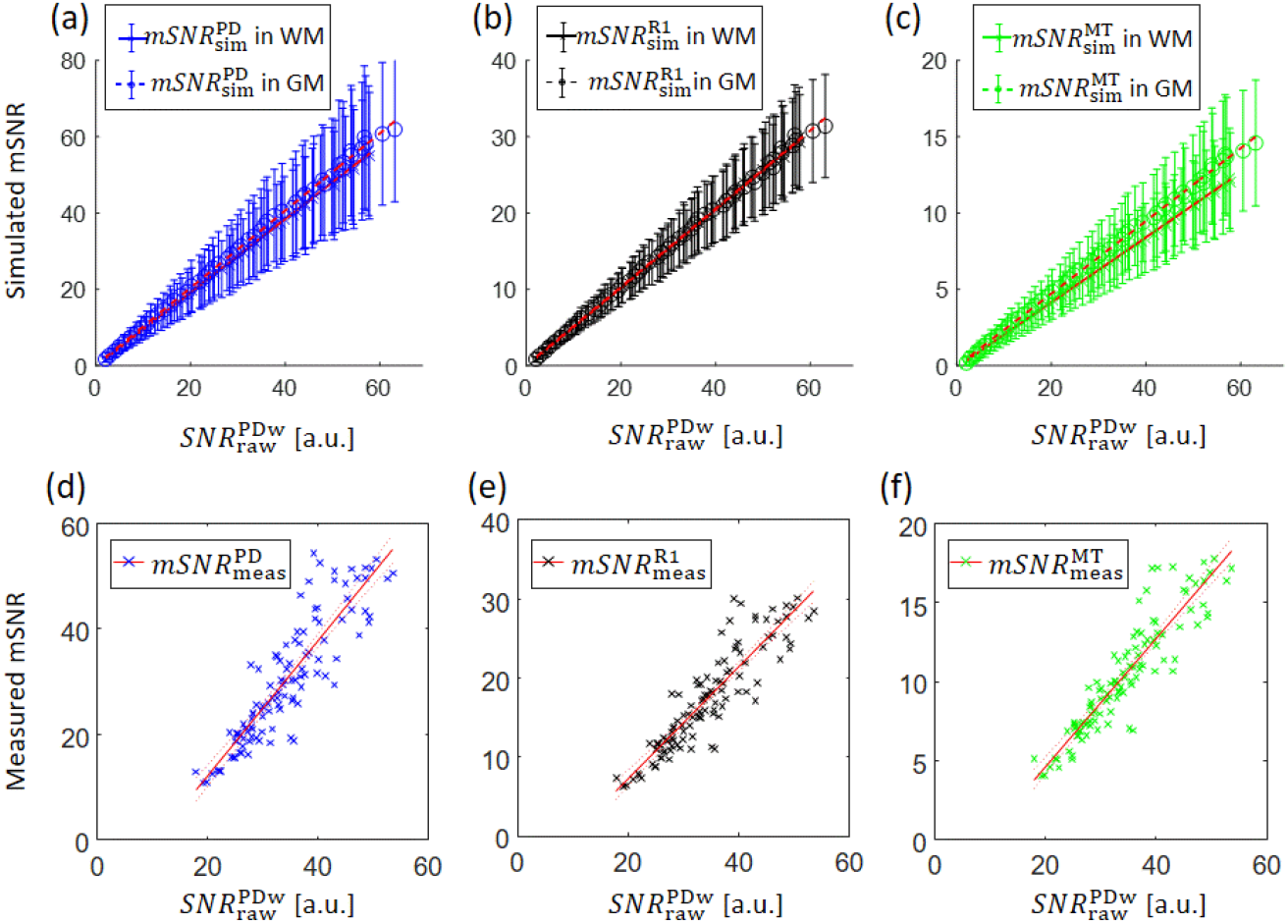
Relating mSNR to the image SNR of the PD-weighted acquisition. Depicted is mSNR as a function of 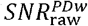 (Eq. (1)) using simulations (a-c) and measurements across a group of healthy subjects (d-f). (a-c): The simulated mSNR is depicted as a function of 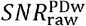 with mean (circle) and standard deviation (errorbar) across 5000 noise realisations for the *PD* (blue), *R*_1_(black), and *MT*_sat_ (green) parameters (Eq. (9)). (c-e): For the ROIs depicted in Figure 9 the mSNR parameters (a: *mSNR*^MT^ as green crosses, b: *mSNR*^R1^ as black crosses, c: *mSNR*^PD^ as blue crosses) are plotted against the corresponding 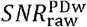 value. A heuristic linear relation is fitted between the mSNR and 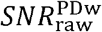 data (Eq. (8)), both for simulated and measured data. The dashed line in magenta is the unity line.

**Figure 9:**
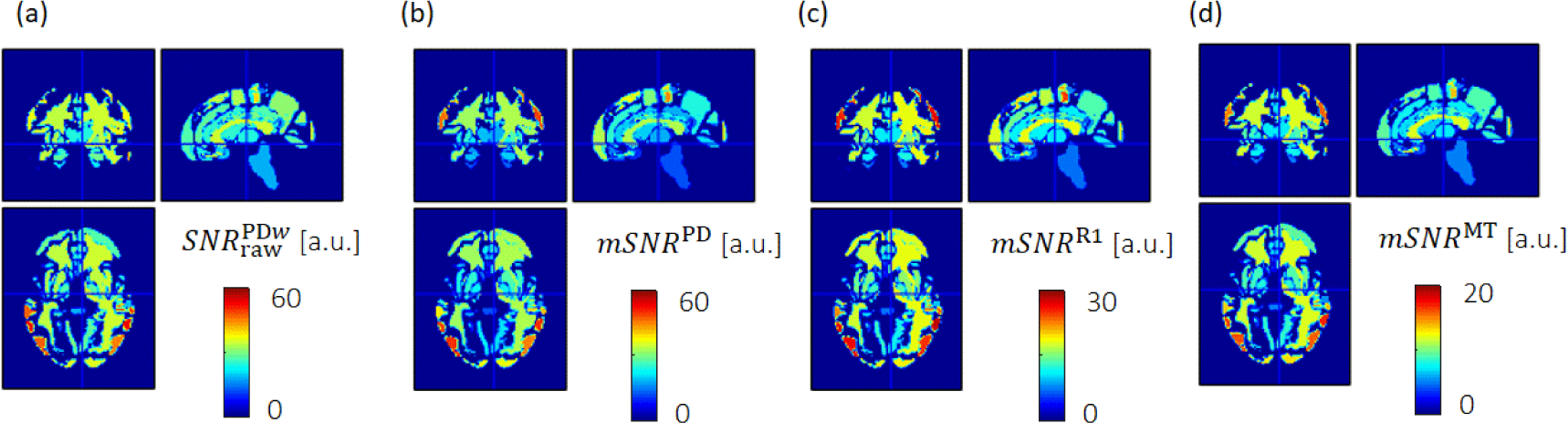
mSNR and 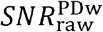 maps averaged within regions-of-interest (ROIs) across the brain. Depicted are the following maps for protocol 1: (a) 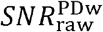, (b) *mSNR*^PD^, (c) *mSNR*^R1^, and (d) *Msnr*^MT^. While the 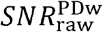 was calculated within each ROI using Eq. (1) per subject, the mSNRs were spatially averaged within each ROI per subject. Then, each of the four different SNR metrics was additionally averaged across the group of healthy subjects on a voxel-by-voxel level. The ROIs were selected using the Oxford-Harvard atlas (Frazier et al., 2005; Desikan et al., 2006; Makris et al., 2006; Goldstein et al., 2007). The 111 ROIs out of 117 were used in which the number of voxels was larger than 100.

The range of each SNR measure was as follows: PD18-54 for 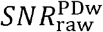,, 11-54 for *mSNR*^PD^, 6-30 for *mSNR*^R1^, and 4-18 for *mSNR*^MT^. We observed a linear relation between 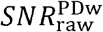 and the mSNRs with highly significant p-values for the slopes but non-significant p-values for the intercept in the simulated data (significance level: p < 0.05 with the null hypothesis in each case being that the parameter is zero). The slope of the mSNR parameter curve was steepest for *mSNR*^PD^, followed by *mSNR*^R1^, and was smallest for *mSNR*^MT^. The simulations revealed that the slope of the mSNR parameter curve was systematically smaller for WM than for GM (2% for *mSNR*^R1^, 6% for *mSNR*^PD^, and 14% for *mSNR*^MT^). We found a similar trend for the ratio of the slopes between measurements and simulations when taking the fitted *mSNR*^PD^-parameter as reference: the slope of *mSNR*^PD^ was 1.8 to 2 times higher than the slope of *mSNR*^R1^, whereas it was 3.2 to 4.5 times higher for *mSNR*^MT^, indicating that MTsat requires a much bigger gain in 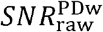 for them to translate into gains in *mSNR*^MT^ as compared to PD. Moreover, we found that the fitted intercept differed substantially between measurements and simulations (Table 3).

### Analysis III: Reducing artefactual variation at the group level

In this analysis, we show how artefactual variations at the group level vary for different combinations of a two-repeat MPM acquisition. Variability at the group level was assessed by the standard-error-of-the-mean (SEM). For WM, the SEM of the arithmetic mean (AM) of the MPM parameters showed opposite variability between *SEM*^PD^ and *SEM*^MT^, e.g.: *SEM*^PD^ was high in the cortical spinal tracts whereas *SEM*^MD^ was low (arrow in Fig. 10a). For GM, the SEM showed higher values toward the outer edge (Fig. 11), potentially caused by residual inaccuracies in spatial registration.

**Figure 10:**
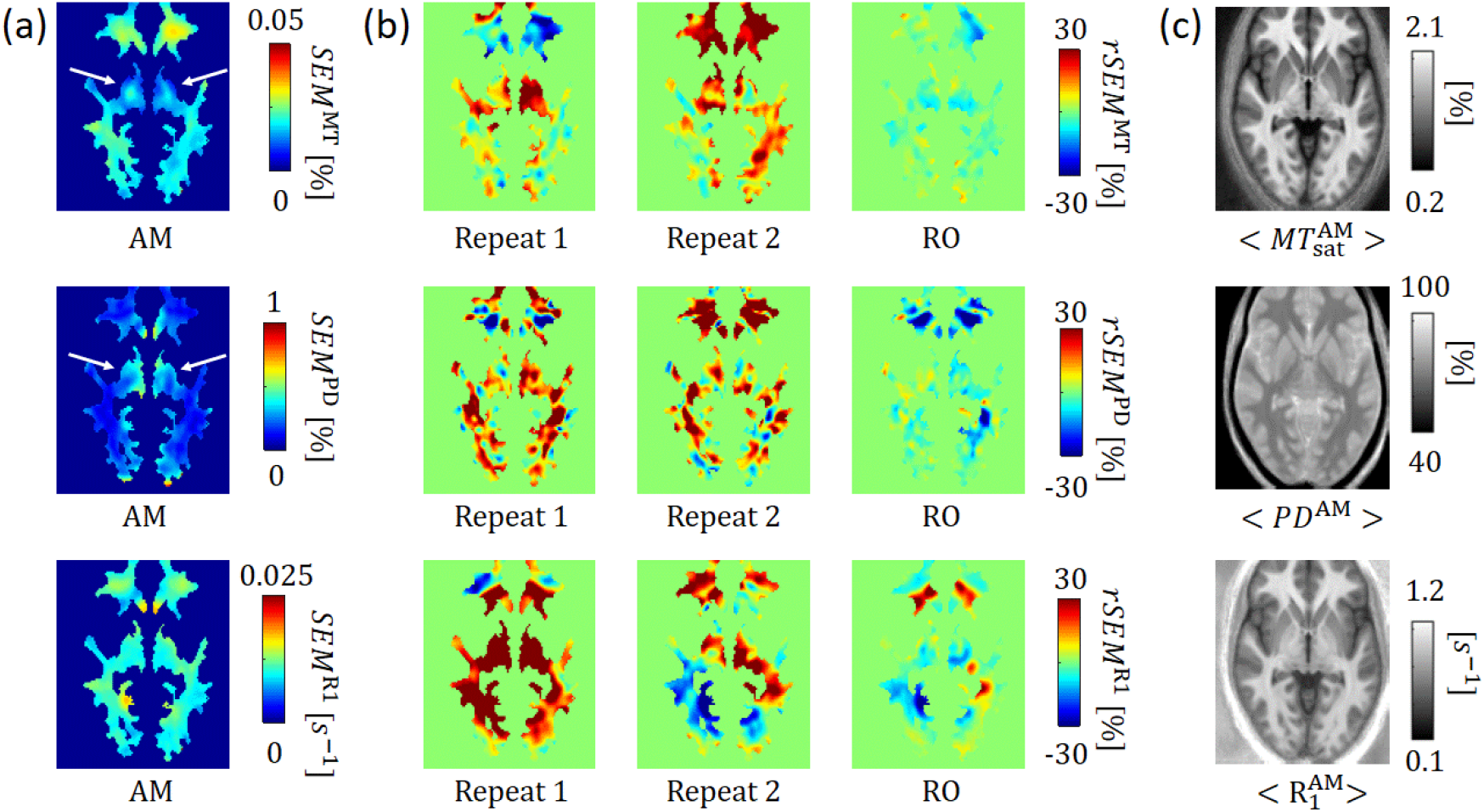
Group variability across subjects for white matter. The group variability was assessed by the standard-error-of-the-mean (*SEM*^*m*^) for the quantitative MPM maps *MT*_sat_ (top row), *PD* (middle row) and *R*_1_ (bottom row) and is illustrated for white matter (*m* ∈ {*MT*, PD, *R*1). Depicted are (a): *SEM*^*m*^ maps generated from the arithmetic mean (AM) of the two-repeat datasets; (b, from left to right): the relative change of *SEM*^*m*^ (denoted as *rSEM*) for the 1^st^ and 2^nd^ repeat dataset, and their robust combination (RO) using *SEM*_*AM*_ as reference; (c): the group-averaged MPM maps. Regions showing reduced SEM are blue and regions showing increased SEM are red. Note that the *SEM*^*m*^ values are not directly comparable, since the scaling of the associated parameters is very different.

**Figure 11:**
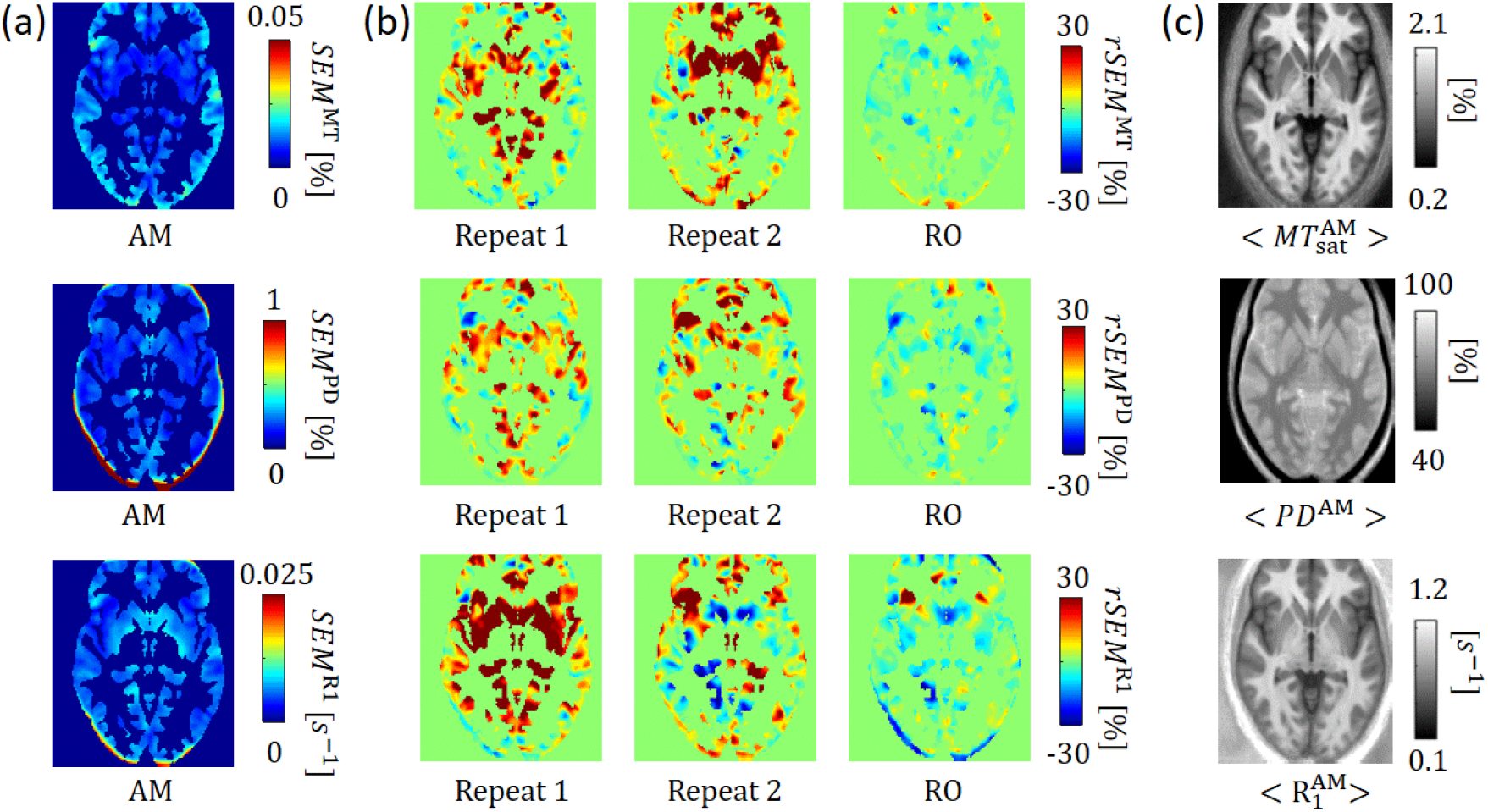
Group variability across subjects for grey matter. Depicted is the same information as in Fig. 10 but for grey matter instead of white matter.

All three MPM parameters showed, on average across the brain, a higher variability if the SEM was estimated on the basis of a single repeat as compared to the AM combined MPM parameters (Table 4): across contrasts the relative SEM for repeat 1 and 2 was between 5.3% (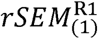 in GM) and 18.7% (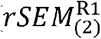 in GM). In some specific regions, however, one of the two repeats showed smaller SEM (blue areas in Figs. 10b and 11b, first and second column). Across the brain, the robust combination (RO) based MPM parameters showed a lower variability as compared to the AM combined MPM parameters: across contrasts the relative SEM was between -0.7 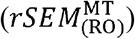 to -4.7% 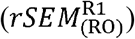. Again, in some localized regions the corresponding r*SEM* maps were also positive, indicating a higher variability after robust combination in specific regions (Figs. 10b and 11b, third column).

**Table 4:** The average relative standard error of the mean (rSEM). The rSEM is estimated with respect to the SEM of the arithmetic-mean (AM) combined MPM parameters 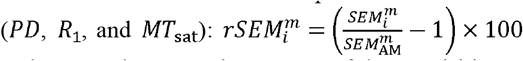 with *m* ∈ {PD,R1,MT} and *i*” ∈ {1,2,RO}, where (1) and (2) are the respective repeats of the acquisition and RO is the robustly combined parameters. Rows 1-3 show the rSEM in white matter (WM) and rows 4-6 show the rSEM in grey matter (GM).

|  | Repeat 1 (1) | Repeat 2 (2) | Robust (RO) |
| --- | --- | --- | --- |
| $rSEM_i^{MT}$ in WM | $7.1 \pm 15.0 \%$ | $11.1 \pm 14.7 \%$ | $-0.8 \pm 4.6 \%$ |
| $rSEM_i^{PD}$ in WM | $14.4 \pm 19.4 \%$ | $9.1 \pm 19.5 \%$ | $-2.9 \pm 9.1 \%$ |
| $rSEM_i^{R1}$ in WM | $14.4 \pm 19.3 \%$ | $9.1 \pm 17.5 \%$ | $-2.9 \pm 8.0 \%$ |
| $rSEM_i^{MT}$ in GM | $10.5 \pm 18.6 \%$ | $12.6 \pm 17.1 \%$ | $-0.7 \pm 6.7 \%$ |
| $rSEM_i^{PD}$ in GM | $9.0 \pm 15.8 \%$ | $6.1 \pm 14.6 \%$ | $-1.0 \pm 8.3 \%$ |
| $rSEM_i^{R1}$ in GM | $18.7 \pm 21.0 \%$ | $5.3 \pm 16.7 \%$ | $-4.7 \pm 10.6 \%$ |

## Discussion

For three quantitative MPM parameters (*R*_1_, *PD, MT*_sat_) we introduced a method to estimate the associated error and model-based signal-to-noise ratio (mSNR) maps without the need to acquire additional data. First, we illustrated that the error and mSNR maps capture the random noise variations associated with instrumental features (e.g. head coil configuration) as well as noise sources related to artefacts (e.g. subject motion). Second, we used measurements across a group of healthy subjects together with simulations to show that mSNRs also reflect SNR. We found that they were linearly related to raw-image-based SNR and that their slopes varied between MPM parameters: the slope was highest for *PD*, lower for *R*_1_ and lowest for *MT*_sat_. Third, we exploited the artefact-sensitivity of the error maps to generate robust MPM parameters from a two-repeat MPM protocol. We showed that artefactual group variability was reduced in the two-repeat MPM acquisition as compared to the single-repeat MPMs. Importantly, the variability was lowest when using the robust MPM parameters as compared to the arithmetic-mean combination of MPM parameters.

### Error and mSNR

To efficiently capture the errors in the MPM parameters (*R*_1_, *PD*, and *MT*_sat_) for each individual MPM experiment, we proposed using error propagation of uncorrelated uncertainties and to approximate the noise variance via the contrast-specific uncertainties of the transverse decay for the PD-, T1-, and MT-weighted SGPR signals (Eq. (4)). The error maps capture noise variation due to random noise and noise associated with artefacts such as subject motion.

The random noise sensitivity is best revealed when using the mSNR which is calculated, in analogy to the SNR, as the ratio between the parameter and its error. The mSNR (Fig. 3a) decreased towards the centre of the brain, which is in accordance with the expected decrease in SNR due to the receive field of the head coil (64ch coil in protocol 1 and 2 and 32ch coil in protocol 3). We found that the change of the mSNR from the brain periphery to the centre was different for protocols 1 and 2 although they were using equivalent MR systems (3T and 64ch head-coil). The most apparent difference in those protocols was the acceleration and the Partial-Fourier (PF) factors: a 3×1 acceleration together with a PF of 6/8 was used in protocol 1 whereas in protocol 2 a 2×2 acceleration and no PF was used. The direction of acceleration in protocol 1 coincided with the direction in which the steeper decline of mSNR values was observed in the respective mSNR maps (left-right direction) as compared to the mSNR acquired with protocol 2. Moreover, the local decrease in mSNR was accompanied by an increase of the contrast-specific uncertainties, meaning that the lower mSNR is driven by a higher noise or artefact level in those regions. One potential reason for the protocol-specific noise-pattern could be an interaction between the g-factor-induced SNR loss (Robson et al., 2008) and Partial Fourier imaging effects. Additionally, the changes in spatial resolution and field strength not only changed the decline of mSNR towards the centre of the brain (protocol 2 showed a steeper decline than protocol 3) but also the averaged mSNR value across the brain.

The noise variations due to artefacts such as subject motion, were better visualized by the error maps but also present in the mSNR. We demonstrated that this second source of variance typically appeared as a local increase in error (and decrease in mSNR) and was accompanied by a bias in the MPM parameters (Fig. 4 and Figs. 5-7).

### Raw-image vs. model-based SNR

We found in simulations and group-level measurements that the mSNR is linearly related to the image-based SNR. This confirms that the mSNR is a genuine measure of SNR. Interestingly, the relative slopes showed a similar trend between simulations and measurements: the slope of *mSNR*^PD^ was 1.8 to 2 times higher than the slope of *mSNR*^R1^, whereas it was 3.2 to 4.5 times higher than for *mSNR*^MT^. The relation between *mSNR*^PD^ and *mSNR*^R1^ will depend on the chosen flip angles and TRs as has previously been shown for SNR in R1 and PD (Helms et al., 2011). A similar argument can be used to understand that *mSNR*^MT^ will also be flip angle dependent.

The following application exemplifies how this information can be relevant for large-scale neuroimaging studies, fundamental neuroscience or clinical research studies. If we assume that two groups of subjects possess different myelin densities in the brain (e.g. due to aging (Callaghan et al., 2014)) and that this difference has the same effect size in *PD* (via *MTV* = 1 – *PD*/100, (Mezer et al., 2013)) and *MT*_sat_ we would need 3.2-4.5 times higher image SNR to observe the same effect under the influence of noise when using *MT*_sat_ as a biomarkersat for myelin instead of *PD*. Only considering noise, this provides a good argument to skip the MT-weighted contrast and thus to further reduce scan time in time-critical studies.

In contrast to the slope of *mSNR*^*m*^, its offset strongly differed between simulations (almost zero) and measurements (−13 for *mSNR*^PT^ MTto *mSNR*^MT^). One reason for this deviation could be that the mSNR in the experiment was compared between different brain regions with different relaxation rates and MTsat values, each of which might possess a slightly different linear dependency between *mSNR*^*m*^ and 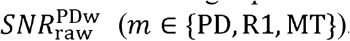. Our simulations revealed that the slope was consistently smaller in WM than in GM (2% for, *R*_1_, 6% for *PD* and 14% for *MT*_sat_). Another reason for the deviation could be the fact that the measured mSNR map is sensitive to both random noise and spatially varying artefacts, e.g., due to parallel imaging, whereas simulations included only random noise.

### Reducing artefactual variation at the group level

The variability at the group level is composed of the true anatomical variability and the artefactual variability induced by different noise sources (e.g. thermal noise, physiological noise, or subject motion). In contrast to the former, the latter type of variability can be reduced by repeated measurements. In accordance with this knowledge, we found that an arithmetic-mean combination of the two repeats reduced the group variability of the MPM parameters (variability reduction: 10.9% in WM and 10.4% in GM) but even more so if the robust combination was used (additional reduction of 2.2% in WM and 2.1% in GM). The additional improvement using robust combination confirms that the reliability of MPM parameters can be further improved when the error map information is exploited to down-weight erroneous MPM values on a per-repeat basis.

Note that we focussed here on the direct effects of the two-repeat protocol on the MPM parameters by using the same transformation and tissue segments for all datasets. However, we believe that the higher artefact level in the single-repeat MPM parameter maps will also degrade the segmentation and by consequence the spatial registration, which, in turn, will further degrade the sensitivity to any true group differences.

### Considerations

The two-repeat protocols are longer in scan time than their one-repeat counterparts (e.g. for protocol 1 it is: 17 min vs. 28 min). Since scan time is often the limiting factor, it is important to consider scan time when comparing variability. To do so, here we translate the reduction in variability into an effective increase in sample size, assessed via the standard-error-of-the-mean (SEM). Under the assumption of Gaussian distributed independent data the SEM directly dictates statistical sensitivity (t-score ∝ 1/SEM) and scales with one-over the square-root of the sample size N. With these relations in mind, a 13% reduction of *SEM*^RO^ relative to the SEM of a standard MPM acquisition (*SEM*^stand^) would translate to an effective increase of the sample size of 31%. This number is directly proportional to the gain in effective scan time, i.e. cumulative scan-time across all subjects. Since a two-repeat acquisition is required for the robust combination, it comes at the price of an extended MPM protocol that was about 39% longer than the corresponding one-repeat acquisition of protocol 1 (total scan time about: 17 min). In total, the effective scan time of the proposed protocol is about 8% longer than the one-repeat protocol, even if accounting for the improved variability due to robust combination. Thus, the proposed protocol and robust combination might be more useful for specific studies, where a small group of subjects, e.g., patients with a rare disease, are investigated and high-quality data of each subject is of higher priority than scan time, but not for studies where a large number of subjects can be afforded (and poor datasets excluded).

In some regions, we found that the proposed robustly combined MPM parameters showed a higher SEM than the arithmetic mean combination, indicating that the error maps do not always correctly capture erroneous regions. One reason might be that the propagation of uncorrelated errors that was used to generate the error maps relied on the assumption that imaging artefacts were adequately captured by the contrast-specific uncertainties estimated from the linearized SPGR fit. This, however, might not always be valid. For example, the contribution from uncertainties in the B1+ estimate is neglected (i.e. we assume 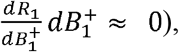 but may play an important role as a source of variance in the MPM parameters (Lee et al., 2017), especially for applications at 7T.

### Potential applications

The knowledge that the mSNR is a genuine measure of SNR can be useful for global and local power analyses. On a local level the mSNR maps might inform studies interested in specific regions regarding which protocol settings, e.g. parallel imaging acceleration or RF head coil, would be optimal. For example, in studies focussing on the hippocampus a less steep decay of mSNR towards the centre of the brain would be preferred, whereas maximal peripheral sensitivity may be preferable for a study interested in the neocortex.

The error and/or mSNR maps can be directly used for statistical comparisons as confidence measures reflecting variation in SNR and erroneous MPM values due to artefact. This allows use of the error maps to improve the robustness of statistical analyses at the group level without needing a two-repeat MPM acquisition. Further research is necessary to find the best statistical neuroimaging framework (e.g. a linear mixed model) that allows integration of confidence maps at the individual subject level (here the error or mSNR maps) with variability measures at the group level (typically standard-error-of-the mean across subjects). Alternatively, error and mSNR maps can be used as additional information in group statistics to assess the reliability of observed differences. For example, if statistical significance in voxel-based statistics between two groups is driven by a few outliers, these might be accompanied by particularly high error values. Thus, the error maps can be used to remove or down-weight erroneous regions in group statistics. This application has been shown to improve statistical significance to detect group differences (David et al., 2017).

## Conclusion

We have introduced a new method to estimate parameter-wise error and model-based SNR (mSNR) maps for three MPM parameters (proton density, PD, longitudinal relaxation rate, R1, and magnetization transfer saturation, *MT*_sat_) on a routine basis without requiring additional data. These new measures can be used to estimate the noise sensitivity of MPM parameters and, if two or more MPM measurements are available, improve their robustness to artefacts such as involuntary subject movement on a per-subject level. The sensitivity to noise might be useful for power-calculations and to compare the suitability of the different MPM parameters as biomarkers in neuroscience or clinical research studies. The improved robustness of MPM parameters might be particularly important in clinical studies where patients with a rare disease are investigated and high data quality is more crucial than high throughput of data. All three advances, the error maps, the mSNR maps, and the robustly combined MPM maps, are available in the open-source hMRI toolbox.

## Acknowledgements

We would like to thank Saskia Helbling and Peter McColgan for sharing the 7T dataset originally published in (McColgan et al., 2021). This project was funded by the ERA-NET NEURON (hMRIofSCI), the Federal Ministry of Education and Research (BMBF; 01EW1711A and B), and the German Research Foundation (DFG Priority Program 2041 “Computational Connectomics”, MO 2397/5□1, DFG Emmy Noether Stipend: MO 2397/4□1), and the Forschungszentrums Medizintechnik Hamburg (fmthh; grant 01fmthh2017). The research leading to these results has received funding from the European Research Council under the European Union’s Seventh Framework Programme (FP7/2007-2013) / ERC grant agreement n° 616905. This project has received funding from the European Union’s Horizon 2020 research and innovation programme under the grant agreement No 681094, and is supported by the Swiss State Secretariat for Education, Research and Innovation (SERI) under contract number 15.0137. SK has been funded by the Max Planck Society, two grants from the German Research Foundation (TRR 169/C8, SFB 936/C7) and the European Union (ERC-2016-StG-Self-Control-677804, Baltic Interreg Programme: Baltic Game Industry). MFC is supported by the MRC and Spinal Research Charity through the ERA-NET Neuron joint call (MR/R000050/1). The Wellcome Centre for Human Neuroimaging is supported by core funding from the Wellcome [203147/Z/16/Z].

## List of Symbols and Acronyms

Symbols: Description
General
hMRI: In vivo histology using MRI
MPM: Multi-parameter mapping
SPGR: SPoiled Gradient Recalled echoes
qMRI: Quantitative MRI
VBM: Voxel-based morphometry
WM: White matter
GM: Grey matter
MPM derivatives
SNR: Signal-to-noise ratio
mSNR: Model-based signal-to-noise ratio
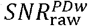: Raw image-based SNR, see Eq. (1)
PD: Proton density
*R*_1_(*T*_1_): Longitudinal relaxation rate (time)
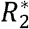: Effective transverse relation rate
*MT*_sat_: Magnetisation transfer saturation rate
*dPD*: Error of *PD*
*dR*1: Error of *R*_1_
*dMT*: Error of *MT*_sat_
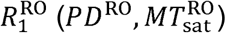: Robustly combined *R*_1_ (PD and MTsat) values from two-repeat acquisition, see Eq. (7)
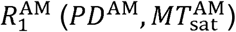: Arithmetic-mean combined *R*_1_ (PD and MTsat) values from two-repeat acquisition
SEM: Standard-error-of-the-mean
rSEM: Relative SEM
Measurement and simulations
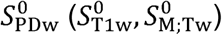: PD-weighted SPGR signal fitted at zero echo time. In brackets: the same for the T1- and MT-weighted SPGR signal
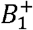: Transmit field
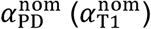: Nominal flip angle for PD (T1)-weighted signal-
*σ*: Standard deviation of noise
*N* (0, *σ*^2^): Zero-mean, additive Gaussian noise with variance *σ*^2^
TE: Echo time
TR: Repetition time
TA: Total acquisition time

